# Interactions between genes altered during cardiotoxicity and neurotoxicity in zebrafish revealed using induced network modules analysis

**DOI:** 10.1101/2022.02.03.478934

**Authors:** Andrea Kagoo R, Ankush Sharma, Anamika Bhargava

**Author notes:** Equal contribution (Andrea Kagoo R, Ankush Sharma).

## Abstract

Zebrafish has become a prominent model organism to study toxicology due to its genomic similarity to humans, optical clarity, well defined developmental stages, short generation time and cost-effective maintenance. It also provides a shorter time frame for *in vivo* toxicology evaluation compared to mammalian experimental systems. As manufacturing processes and development of new synthetic compounds increase to keep pace with the expanding global demand, environmental health and the effects of toxicant exposure are emerging as critical public health concerns. Owing to this, in this study, we analyzed the impact of such chemical induced toxicity in zebrafish. We first searched the literature comprehensively for genes that were modulated during neurotoxicity and cardiotoxicity in zebrafish and then studied the interactions between those genes using induced network modules analysis of the database ConsensusPathDB. The induced network modules analysis helps to study gene interactions through various types of interactions like gene regulatory, biochemical, genetic and protein interactions. A constant communication between the heart and the brain is an important physiological phenomenon. Therefore, we also analyzed the interactions among genes that are modulated simultaneously during cardiotoxicity and neurotoxicity. This study has led us to identify some potential predictive biomarkers for neurotoxicity and cardiotoxicity in zebrafish.

## 1. INTRODUCTION

Zebrafish (*Danio rerio*) is a now a well-known vertebrate model organism to study toxicology [1]. Advantages like cost-effective maintenance, high fecundity, external fertilisation, short life cycle, fast growth rate, well-defined developmental stages, transparency of zebrafish embryos and small size of the embryos as well as adults has made it ideal for toxicity testing. Also, zebrafish has a lot a physiological and functional similarities with humans [2–4]. Approximately 70% of human genes have functional orthologs in the zebrafish [5]. Further, no ethical approval is required to study zebrafish larvae up to 120 hours post fertilisation (hpf) as they are not capable of independent feeding and depend entirely on the yolk supply [6]. The use of zebrafish larvae also brings direct benefits of the 3R’s principle (Replace, Reduce, Refine). They **replace** higher order animals like mice and rats. These complex lower vertebrate animals that emerge from the eggs 48 hpf with functional central nervous system, endocrine and gastro-intestinal and cardiovascular systems allow simultaneous monitoring of behaviour and various physiological parameters. They are also comparatively easier to handle due to small size. Use of zebrafish **reduces** the overall use of mice and rats. It also reduces the cost, chemicals and biowaste produced. It is easier to maintain zebrafish in the natural conditions than is possible to maintain for mammals. This **refines** the animal usage and reduces the housing stress and the impact of that stress on the outcome [7].

In this study, we determined interactions between genes altered during induced cardiotoxicity and neurotoxicity reported in the zebrafish using the database ConsensusPathDB [8]. Neurotoxicity is defined as “an adverse change in the structure or function of the nervous system that results from exposure to a chemical, biological or physical agent” [9]. Advantages like easy penetration of chemicals through the external membrane of zebrafish by passive diffusion, development of the blood brain barrier similar to mammals and fast brain development has made zebrafish ideal for neurotoxicity screening [10]. Neurotoxicity endpoints include gene expression patterns, neural morphogenesis and neuro-behavioural profiling [10]. For the analysis in this study, we considered altered brain gene expression as neurotoxicity. Cardiotoxicity, on the other hand, is defined as the toxicity that damages the heart muscle and other cardiac tissues and/or disrupts the electrophysiology of the heart [11]. The heart rate and action potential of zebrafish are analogous to humans [12]. Along with this, transparency allowing non-invasive techniques and whole animal imaging has made zebrafish ideal for cardiotoxicity evaluation [13]. Cardiac function assessment can be carried out by investigating the teratogenic effects and by a variety of hemodynamic parameters like heartbeat, cardiac output, fractional area change, fractional shortening, and vascular blood flow velocities and altered gene expression. For the analysis in this study, we considered altered cardiac gene expression as cardiotoxicity. Owing to rising critical health concerns due to toxicant exposure, this study focuses on the impact of such chemicals on gene expression in neurotoxicity and cardiotoxicity in zebrafish. It also aims to collate otherwise scattered information of organ-specific modulators for neurotoxicity and cardiotoxicity that will help to understand the mechanisms of toxicity by studying the interaction networks between them. ConsensusPathDB integrates different types of functional interactions between physical entities in the cell like genes, RNA and proteins [14].

Here, we first manually curated the articles related to neurotoxicity and cardiotoxicity in zebrafish published in 2019 and 2020. We then subjected the genes altered during neurotoxicity and cardiotoxicity with a fold change greater than 3 to induced network modules analysis individually. Pathophysiological interplays between the nervous and cardiovascular systems prompted us to investigate if there were any common target genes between neurotoxicity and cardiotoxicity and the interactions between them.

## 2. METHODS

We searched the literature for articles that reported cardiotoxicity and neurotoxicity in zebrafish in the year 2019 and 2020 using search engines PubMed and google scholar. We found 592 articles for neurotoxicity and 218 articles for cardiotoxicity. Then the articles were manually screened for gene expression studies and data such as gene name, gene alteration (upregulated or downregulated), primer sequences, study design including the chemical used, time of exposure, time point at which qPCR was done, strain and the developmental stage of the zebrafish. Toxicity articles that did not report gene expression data were excluded from our analysis.

A total of 76 articles for neurotoxicity and 34 articles for cardiotoxicity were found relevant and were included in the analysis (Titles of these articles are provided in the supplementary information table S1 and S2 respectively). From the data collected from the relevant articles, the gene lists were created and divided based on the alteration of expression observed. Six gene lists were created: 1) Exclusively upregulated genes in neurotoxicity (supplementary information table S3) 2) Common upregulated genes in neurotoxicity and cardiotoxicity (supplementary information table S4) 3) Exclusively downregulated genes in neurotoxicity (supplementary information table S5) 4) Common downregulated genes in neurotoxicity and cardiotoxicity (supplementary information table S6) 5) Exclusively upregulated genes in cardiotoxicity (supplementary information table S7) and 6) Exclusively downregulated genes in cardiotoxicity (supplementary information table S8). Table 1 shows the number of genes in each list. From these gene lists only those genes having fold change greater than 3 were considered for induced network modules analysis to induce stringency in interaction analysis. The stringent gene lists were then submitted into the Induced network modules analysis section under gene set analysis in the **ConsensusPathDB** tool and analysed.

**Table 1:** Numbers of genes in each list.

| **List** | **Total genes** | **Genes with fold change greater than 3** |
| --- | --- | --- |
| Exclusively upregulated genes in neurotoxicity | **59** | **19** |
| Common upregulated genes in neurotoxicity and cardiotoxicity | **18** | **6** |
| Exclusively downregulated genes in neurotoxicity | **118** | **40** |
| Common downregulated genes in neurotoxicity and cardiotoxicity | **10** | **2** |
| Exclusively upregulated genes in cardiotoxicity | **14** | **6** |
| Exclusively downregulated genes in cardiotoxicity | **49** | **22** |

## 3. RESULTS AND DISCUSSION

### 3.1 Interaction between genes upregulated exclusively during neurotoxicity

Since we analysed all genes that were reported to be modulated during neurotoxicity and cardiotoxicity, first we wanted to understand the interaction between genes that were exclusively upregulated during neurotoxicity. In this, genes that were upregulated during neurotoxicity as well as cardiotoxicity were excluded.

#### Gene regulatory interactions

The list of genes was subjected to gene regulatory interactions in the **Induced network modules analysis** in ConsensusPathDB. Gene regulatory interactions provide a map or blueprint of molecular interactions. Such networks can help to derive a novel hypothesis which then can be investigated experimentally. These networks can also be used as biomarkers for diagnostic, prognostic or predictive purposes [15]. The analysis yielded three clusters (Fig. 1). Seed nodes are genes in our input list and intermediate nodes are likely to be associated with the phenotype under study, although they may not be regulated on the transcriptional level and thus do not appear in the input gene list [14]. The first cluster comprised of seed node IL10 which is a cytokine with potent anti-inflammatory properties [16]. Intermediate nodes included transcription factors like SMAD3/SMAD4/GATA3 and JUN/JUNB. The second cluster consisted of seed nodes like NFE2L2, HMOX1 and CXCL8. Nuclear Factor, Erythroid 2 Like (NFE2L2) is a transcriptional activator that binds to the antioxidant response elements (ARE) and protects against oxidative stress by upregulating other antioxidant proteins [17]. Heme oxygenase (HMOX1), an essential enzyme in heme catabolism, cleaves heme to form biliverdin, which is subsequently converted to bilirubin by biliverdin reductase, and carbon monoxide, a putative neurotransmitter [18]. This gives it an anti-inflammatory property by upregulating IL10 [19]. It also exhibits cytoprotective effects since excess of free heme sensitizes cells to undergo apoptosis [20]. C-X-C Motif Chemokine Ligand 8 (CXCL8) (interleukin-8) is a major mediator of the inflammatory response[21]. Intermediate nodes like CK2 complex, BLVRA (Biliverdin reductase A), Fra1/USF2, p-T180, Y182-MAPK14, Inhibitor of nuclear factor kappa B kinase beta subunit (IKBKB), HB-EGF/EGFR, ATF2/JUND/macroH2A and Fra1/JUND were found. The third cluster was formed by seed nodes like PRKAB1, tumor suppressor p53, ornithine decarboxylase and intermediate node MYC/Max/Cbp/p300.

**Figure 1.**
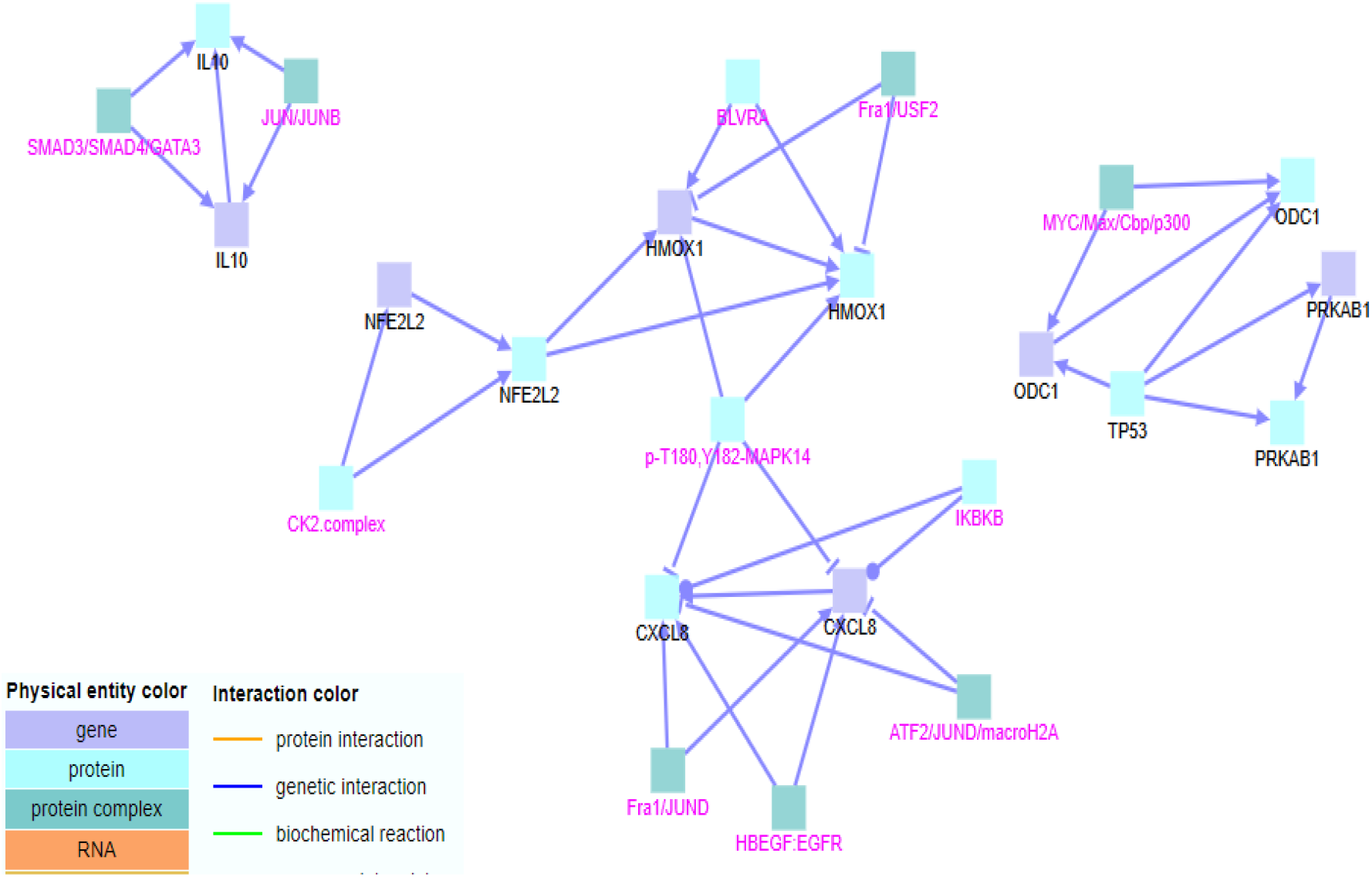
Gene regulatory interactions between genes exclusively upregulated in neurotoxicity. Black labels denote seed nodes and magenta labels denote intermediate nodes. Each edge represents an interaction.

#### Biochemical reactions

When the same list was subjected additionally to biochemical interactions, all the clusters interacted (Fig. 2). Additional seed nodes LTA (Lymphotoxin alpha) and TGFB3 (Transforming Growth Factor Beta 3) were included. LTA mediates a large variety of inflammatory and immunostimulatory responses [22]. Additional intermediate nodes like HIF-1 complex, HIF1A/ARNT, NQO1, MAF, SP3, NFkB complex, ATF1, p-S407, Y641-STAT6 dimer, PBX1, Fra1/JUN, p300/CBP/RELA/p50, HIST3H3, JUN/FOS/NFAT1-c-4 and p-2S-cJUN: p-2S,2T-cFOS were found. Hypoxia-Inducible Factor (HIF)-1 is a dimeric protein complex that plays an essential role in the body’s response to low oxygen concentrations (hypoxia) [23]. NQO1 serves as a quinone reductase in connection with conjugation reactions of hydroquinones involved in detoxification pathways [24]. NFkB plays a major role in the regulation of inflammatory responses [25]

**Figure 2.**
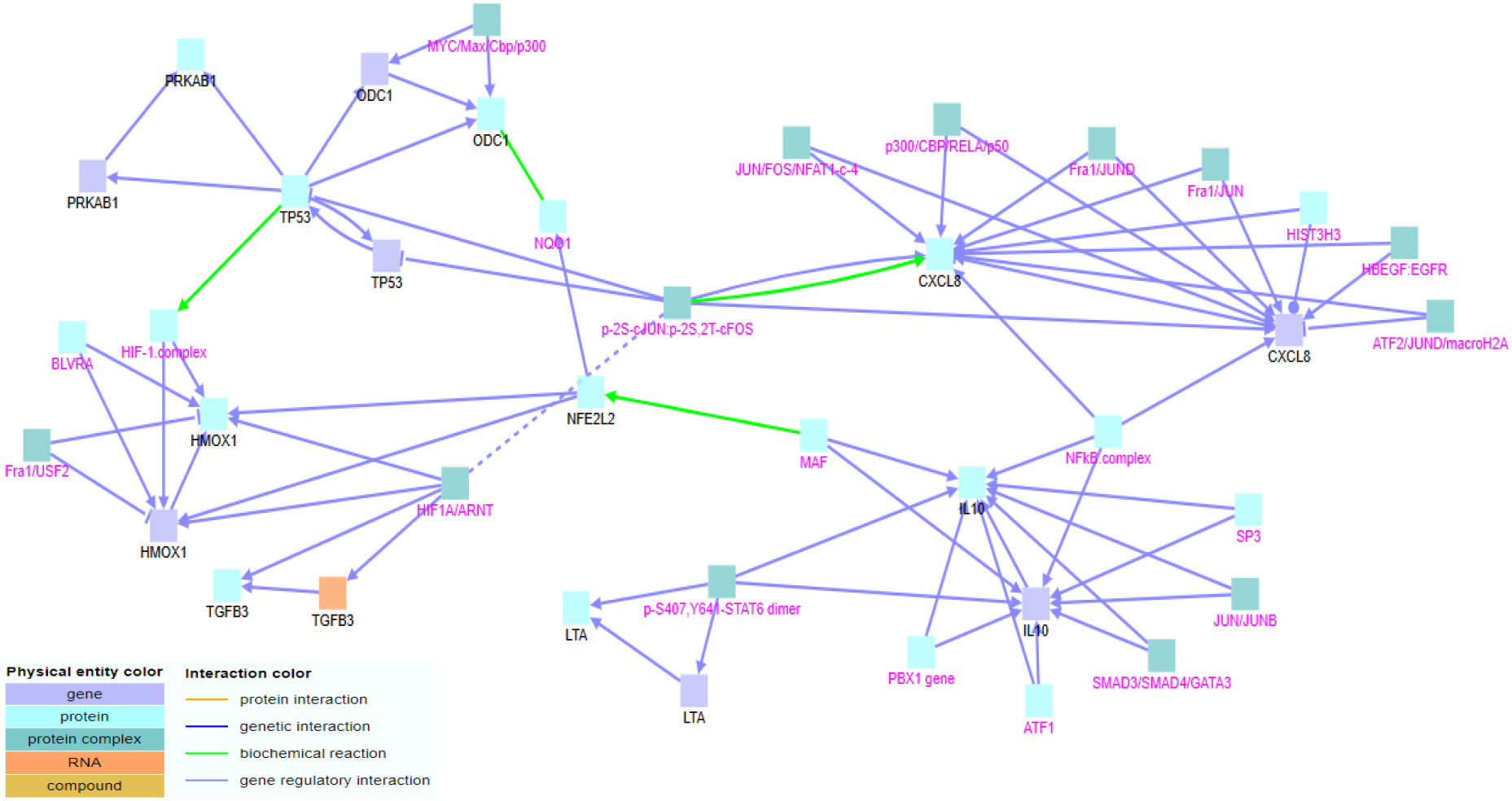
Gene regulatory interactions and biochemical reactions between genes exclusively upregulated in neurotoxicity. Black labels denote seed nodes and magenta labels denote intermediate nodes. Each edge represents an interaction.

#### Genetic interactions

Genetic interactions help us to understand the mechanisms underlying the robustness of biological systems. It gives a useful understanding of the relationship between the genotype and phenotype [26]. When genetic interactions were included in the analysis, an interaction was found between p53 gene and CXCL8 (Fig. 3). p53 is known to be involved in the regulation of several proinflammatory genes in human macrophages like IL-6, IL-8 (CXCL8) and CXCL1 [27].

**Figure 3.**
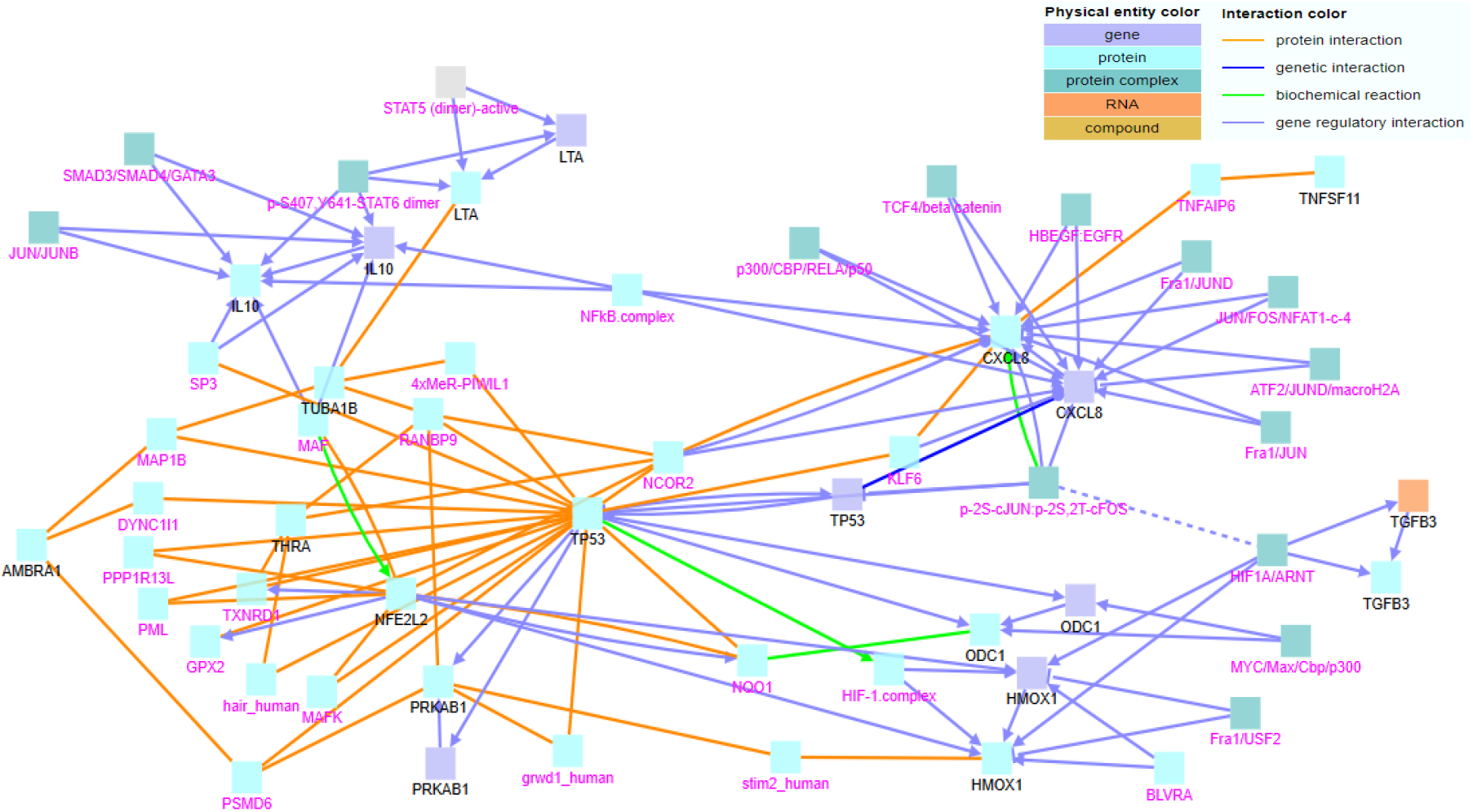
Gene regulatory, biochemical reactions, genetic and protein interactions between genes exclusively upregulated in neurotoxicity. Black labels denote seed nodes and magenta labels denote intermediate nodes. Each edge represents an interaction.

#### Protein interactions

Protein interactions in networks are increasingly used to decipher the molecular basis of disease states [28]. When protein interactions were included in the analysis, seed nodes like AMBRA1, TUBA1B, THRA and TNFSF11 were included in the cluster (Fig. 3). Additional intermediate nodes like TNFAIP6, TCF4/beta catenin, MAFK, STAT5 (dimer)-active, KLF6, NCOR2, STIM2, RANBP9, Piwi like 1, PPP1R13L, DYNC1I1, GPX2, PML, PSMD6, TXNRD1, Hairless Protein, MAP1B, GRWD1 were found. GPX2 (Glutathione Peroxidase 2) catalyze the reduction of organic hydroperoxides and hydrogen peroxide (H_2_O_2_) by glutathione and hence protects cells against oxidative damage [29]. MAP1B (Microtubule Associated Protein 1B) is known to be involved in microtubule assembly, which is an essential step in neurogenesis[30]. GRWD1 (Glutamate-rich WD repeat-containing protein 1) is a histone binding-protein that regulates chromatin dynamics [31]. Hairless protein is a potent antagonist of neurogenic gene activity during sensory organ development[32]. KLF6 is a zinc finger protein that is a transcriptional activator, and functions as a tumor suppressor [33].

### 3.2 Interactions between genes upregulated both in neurotoxicity and cardiotoxicity

#### Gene regulatory interactions

When the list comprising common upregulated genes in neurotoxicity and cardiotoxicity was subjected to gene regulatory interactions in ConsensusPathDB, one cluster was obtained (Fig. 4). Seed node Prostaglandin G/H synthase 2 precursor (PTGS2) was included along with intermediate nodes p38alpha-beta-active, ERBB2, JNK family, Patatin-like phospholipase domain-containing protein 8 (PNPLA8), phospholipase A2 group VI (PLA2G6), JUN family and ERK family. ERBB2 is a member of the epidermal growth factor receptor (EGFR) family of receptor tyrosine kinases. Overexpression of this gene has been reported in numerous cancers [34].

**Figure 4.**
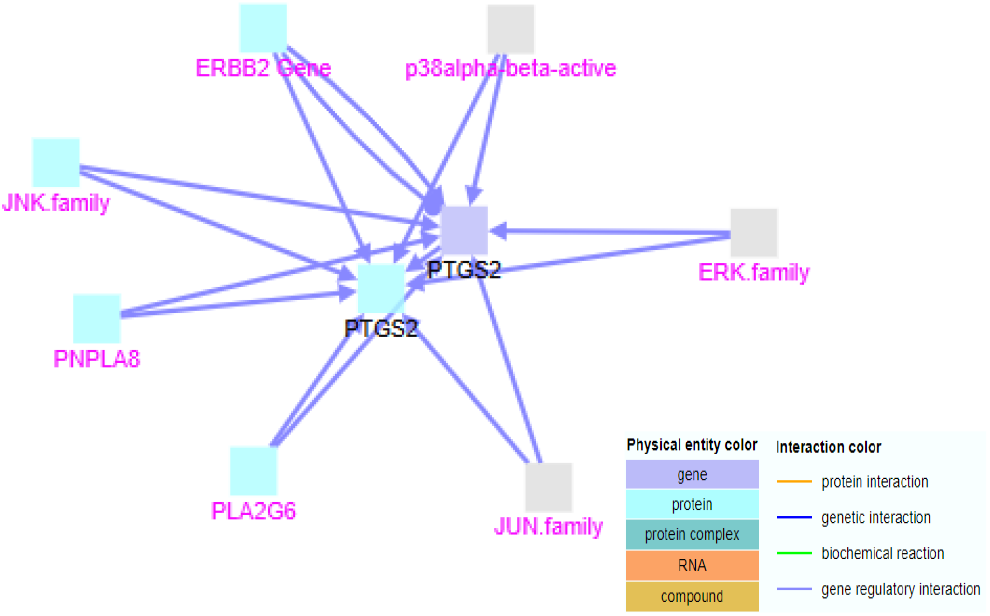
Gene regulatory interactions between genes upregulated both in neurotoxicity and cardiotoxicity. Black labels denote seed nodes and magenta labels denote intermediate nodes. Each edge represents an interaction.

#### Biochemical reactions

When biochemical reactions were included in the analysis, additional seed nodes like Cytochrome P450 1A1 (CYP1A1) and Homeodomain interacting protein kinase 2 (HIPK2) were included (Fig. 5). Induction of CYP1A1 expression and activity serves as a biomarker of toxicity in diverse animal species and humans [35]. HIPK2 activates certain transcription factors like CREB1 and ATF1 by phosphorylation in response to genotoxic stress [36]. Additional intermediate nodes like p-S63, S73-JUN, Peroxisome proliferator-activated receptor gamma (PPARG), CCAAT/enhancer-binding protein beta (CEBPB), Forkhead box protein L2 (FOXL2), AP-1_I.complex and JUN/FOS/NFAT1-c-4 were found. The protein encoded by the intron less CEBPB gene is important in the regulation of genes involved in immune and inflammatory responses and has been shown to bind to the IL-1 response element in the IL-6 gene [37].

**Figure 5.**
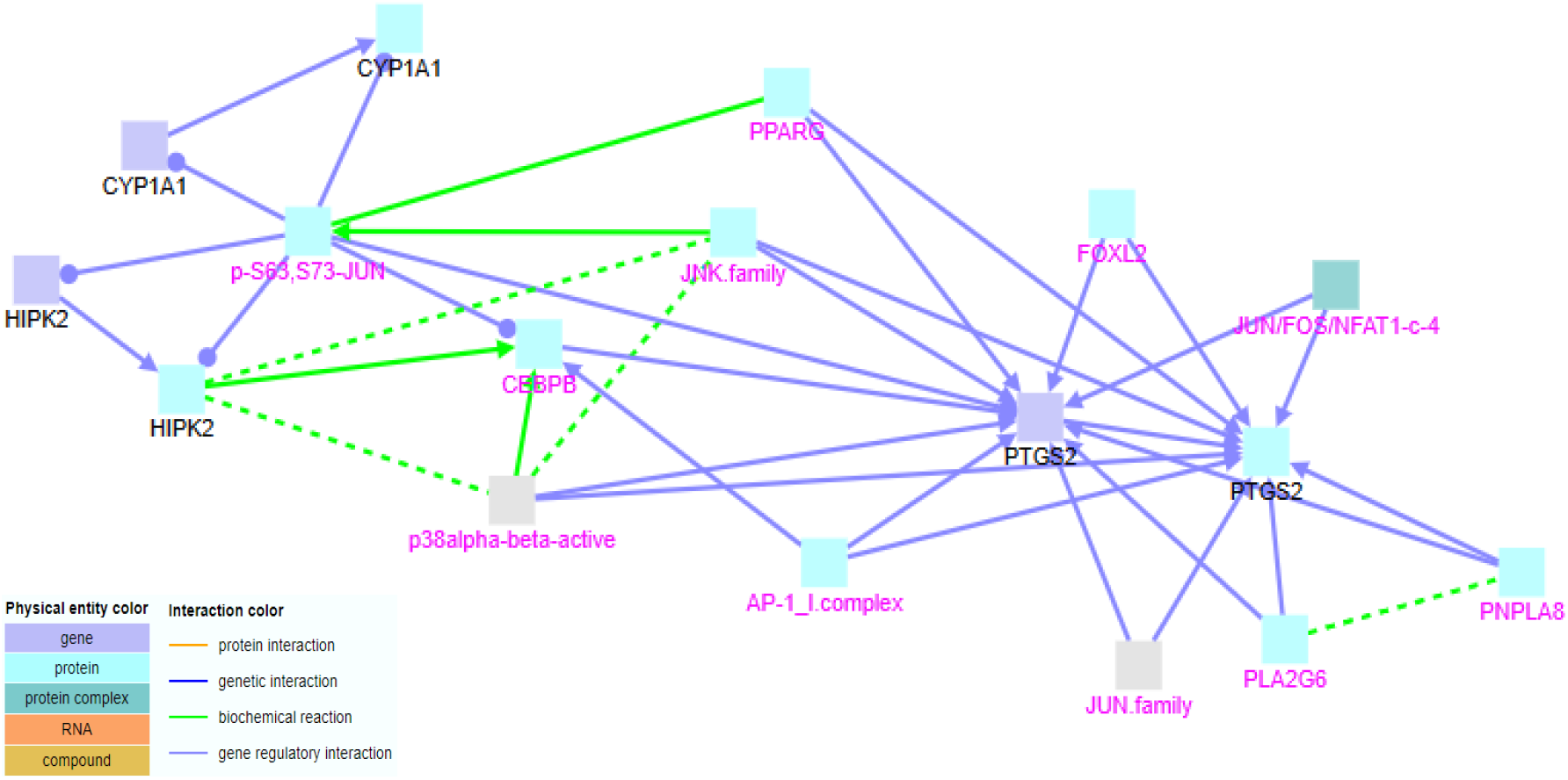
Gene regulatory interactions and biochemical reactions between genes upregulated both in neurotoxicity and cardiotoxicity. Black labels denote seed nodes and magenta labels denote intermediate nodes. Each edge represents an interaction. No genetic interactions were found.

#### Protein interactions

When protein interactions were included in the analysis, additional seed nodes like Interleukin-21 receptor (IL21R) and Aquaporin-1 (AQP1) and additional intermediate nodes like Beta-actin-like protein 2 (ACTBL2), Nucleobindin-1 (NUCB1), COP9 signalosome complex subunit 3 (COPS3), Myeloblastosis viral oncogene homolog (MYB) and 26S proteasome non-ATPase regulatory subunit 1 (PSMD1) were included (Fig. 6). NUCB1 is a major calcium-binding protein of the Golgi which may have a role in calcium homeostasis [38]. MYB protein plays an essential role in the regulation of haematopoiesis. This gene may be aberrantly expressed or rearranged or undergo translocation in leukemias and lymphomas. Also considered to be an oncogene [39]. PSMD1 complex plays a key role in the maintenance of protein homeostasis by removing misfolded or damaged proteins, which could impair cellular functions, and by removing proteins whose functions are no longer required. Therefore, the proteasome participates in numerous cellular processes, including cell cycle progression, apoptosis, or DNA damage repair [40].

**Figure 6.**
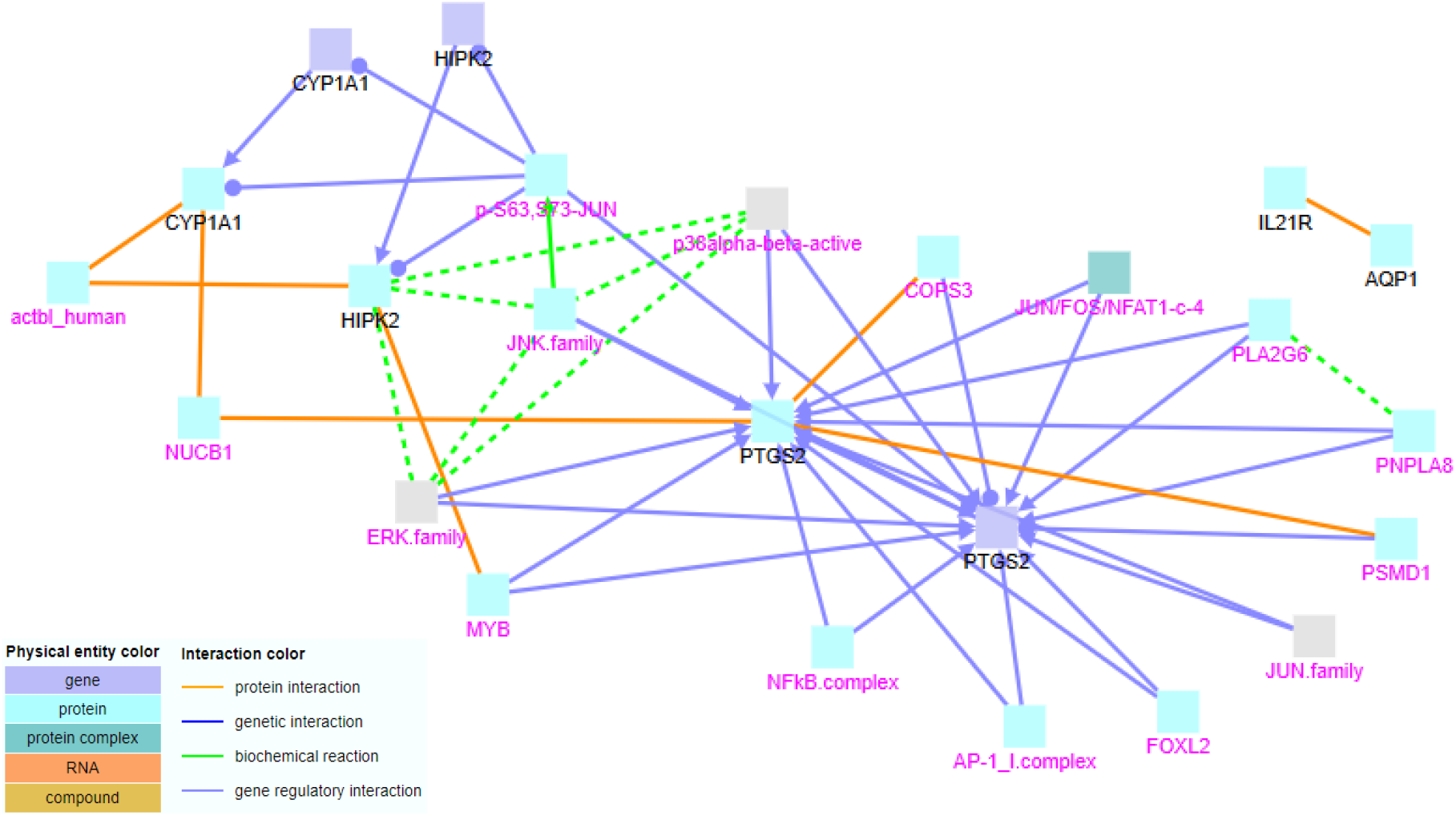
Gene regulatory, biochemical reactions and protein interactions between gene upregulated both in neurotoxicity and cardiotoxicity. Black labels denote seed nodes and magenta labels denote intermediate nodes. Each edge represents an interaction.

### 3.3 Interactions between genes downregulated exclusively in neurotoxicity

#### Gene regulatory interactions

When the list containing the downregulated genes in neurotoxicity excluding the common downregulated genes in neurotoxicity and cardiotoxicity, was subjected to gene regulatory interactions, five clusters were formed (Fig. 7). The first cluster comprised of seed node Glutamate receptor 2 (GRIA2) and intermediate node FOSB: JUND. GRIA2 is a receptor for glutamate that functions as ligand-gated ion channel in the central nervous system and plays an important role in excitatory synaptic transmission. In the presence of CACNG4 or CACNG7 or CACNG8, shows re-sensitization which is characterized by a delayed accumulation of current flux upon continued application of glutamate [41]. The second cluster consisted of seed node Estrogen receptor alpha (ESR1) and intermediate nodes ATF2/JUN/ER alpha, ATF2 (dimer)/ER alpha, JUN/FOS/ER alpha and MPG. The third cluster consisted of seed nodes Suppressor of cytokine signalling 3 (SOCS3) and Phosphatidylinositol 3-kinase catalytic subunit, alpha isoform (PIK3CA). Intermediate nodes like IL2/IL2R alpha/beta/gamma/JAK1/LCK/JAK3, STATS/STATS, p38alpha-beta-active, Jak2/Leptin Receptor and MIR203 were found. The fourth cluster consists of seed node FOS and intermediate nodes like TCF complex, SAP1 (Serum response factor accessory protein 1), ERK1-2/ELK1 and RHOA/GTP. The fifth cluster consisted of seed node Matrix metalloproteinase-9 (MMP9) and intermediate nodes JUNB/FOSB, Fra1/FIAT and FGFR-FGF2 complex/N-cadherin.

**Figure 7.**
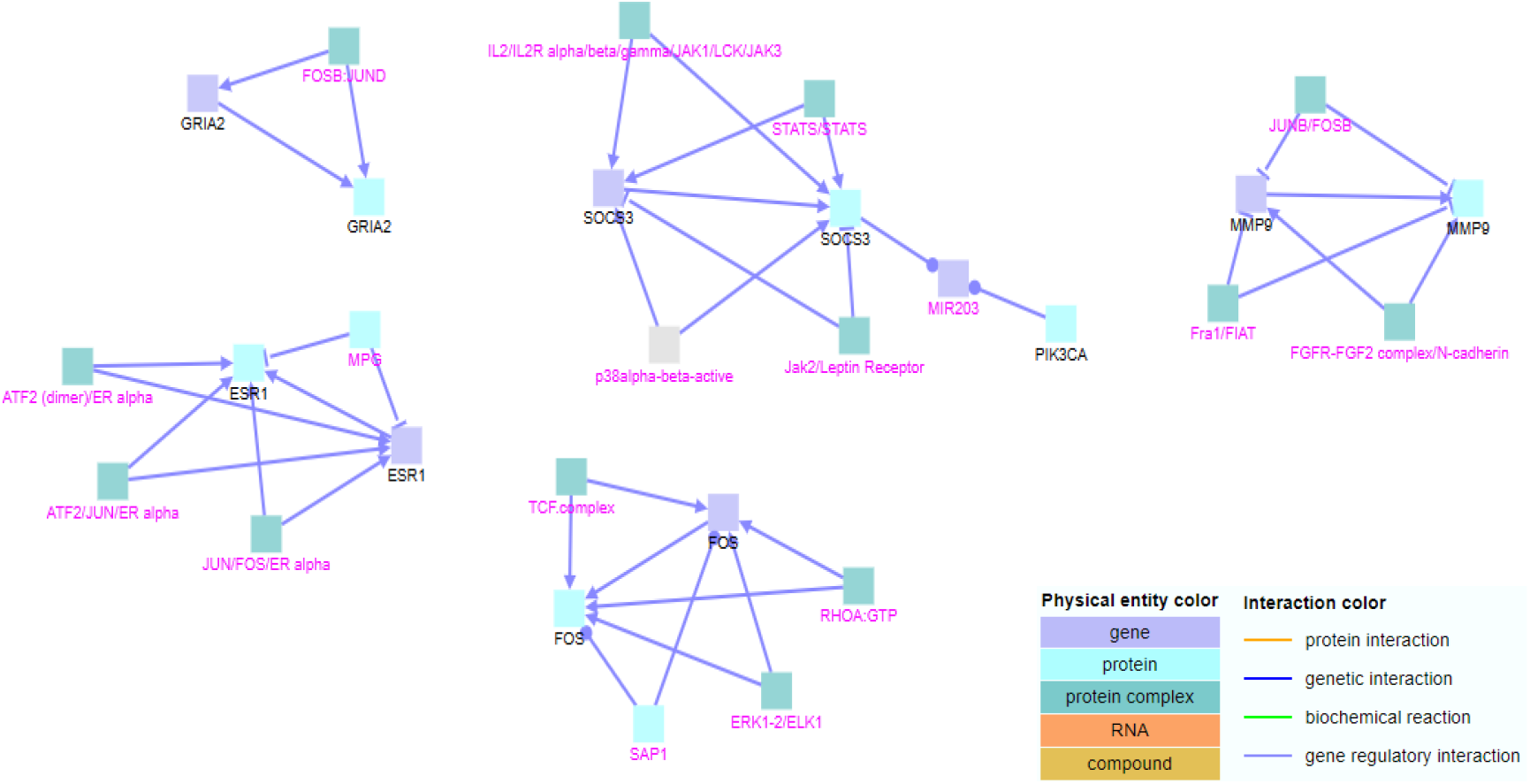
Gene regulatory interactions between genes downregulated exclusively in neurotoxicity. Black labels denote seed nodes and magenta labels denote intermediate nodes. Each edge represents an interaction.

#### Biochemical reactions

When biochemical reactions were included in the analysis, all the clusters interacted (Fig. 8). Additional seed nodes like MTOR, CDK6, CDC42, CACNA1D, CACNA1F, CACNA1A, CACNA1I, EGR2, ARRB2 and LHCGR were found. The protein encoded by the Mammalian target of rapamycin (mTOR) gene belongs to a family of phosphatidylinositol kinase-related kinases. These kinases mediate cellular responses to stresses such as DNA damage and nutrient deprivation. This protein acts as the target for the cell-cycle arrest and immunosuppressive effects of the FKBP12-rapamycin complex [42]. The protein encoded by the CDK6 gene is a member of the cyclin-dependent protein kinase (CDK) family and are important regulators of cell cycle progression [43]. They are known to be over-expressed in some leukaemia and malignancies (including sarcoma, glioma, breast tumours, lymphoma and melanoma) [44]. Voltage gated calcium channels like CACNA1D, CACNA1F, CACNA1A, CACNA1I (the first three being L-type channels and the latter being T-type) can be targets of a number of naturally occurring toxins, therapeutic agents as well as environmental toxicants [45]. Beta-Arrestin 2, isoform 1 (ARRB2) is a major regulator of GPCR signalling by binding to the activated GPCR and causing receptor desensitization and internalization. It is also involved in linking GPCRs to clathrin-coated pits, regulation of cytoskeletal rearrangement and cellular localization, translocation, or stability regulation of signalling elements[46]. Mutations in the luteinizing hormone/choriogonadotropin receptor (LHCGR) result in disorders of male secondary sexual character development, such as familial male precocious puberty and male pseudohermaphroditism [47]. A genetic interaction was found between seed node ESR1 and intermediate node TFF1 gene (Fig 9).

**Figure 8.**
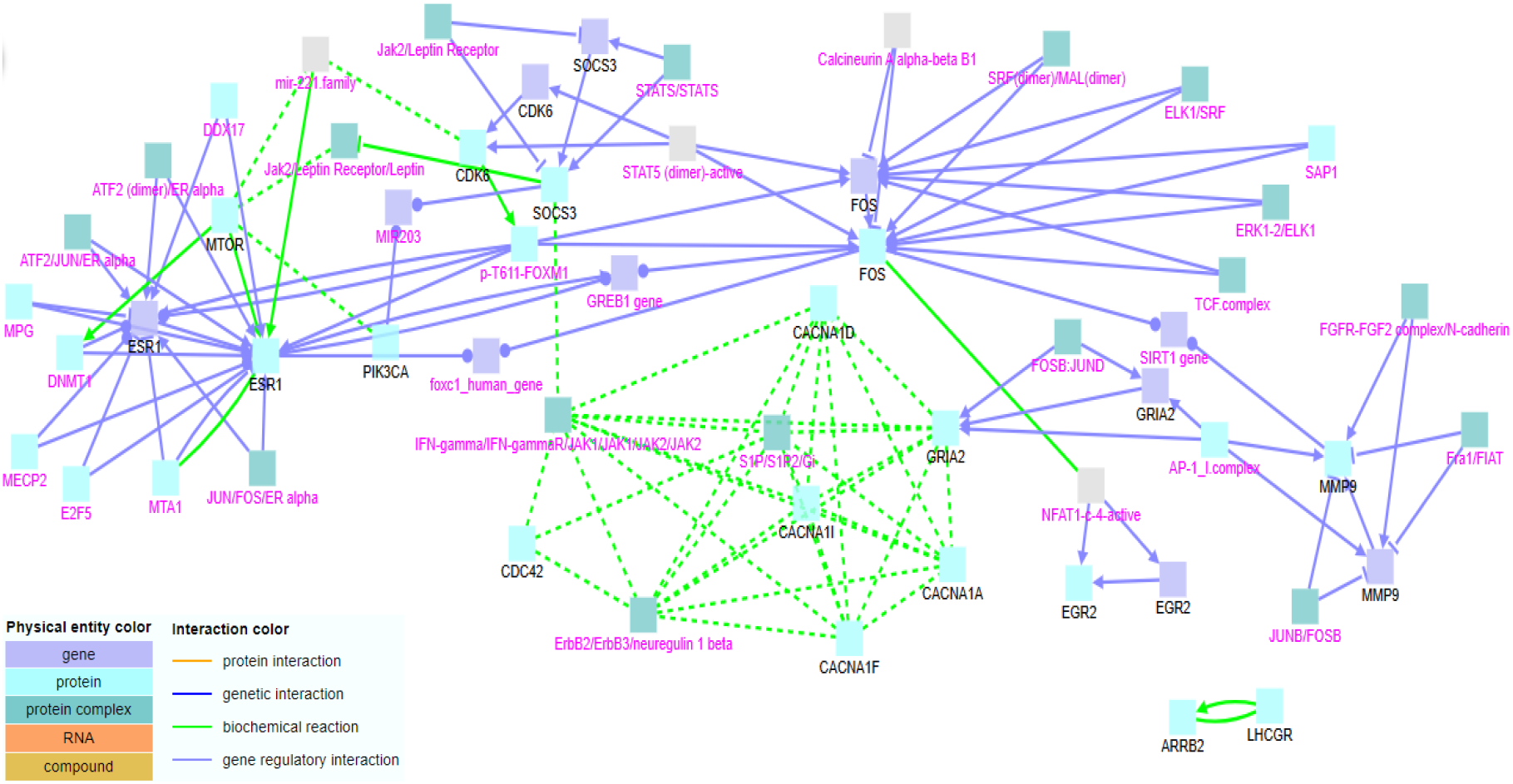
Gene regulatory interactions and biochemical reactions between genes downregulated exclusively in neurotoxicity. Black labels denote seed nodes and magenta labels denote intermediate nodes. Each edge represents an interaction.

**Figure 9.**
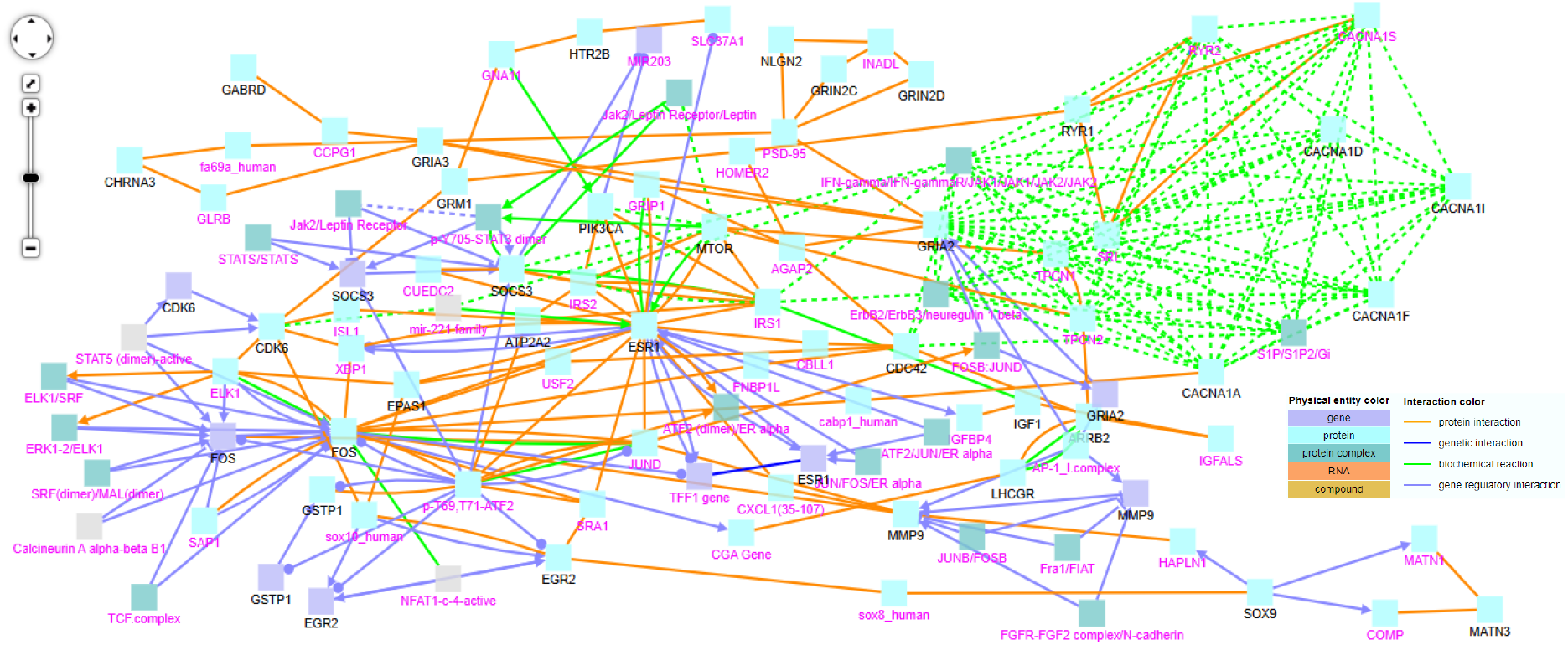
Gene regulatory, biochemical reactions, genetic and protein interactions between genes downregulated exclusively in neurotoxicity. Black labels denote seed nodes and magenta labels denote intermediate nodes. Each edge represents an interaction.

#### Protein interactions

When protein interactions were included, additional seed nodes like GABRD, CHRNA3, GRIA3, GRM1, GSTP1, EPAS1, ATP2A2, HTR2B, SOX9, MATN3, NLGN2, GRIN2C, GRIN2D, IGF1 and RYR1 were included (Fig. 9). Mutations in the gamma-aminobutyric acid type A receptor delta subunit (GABRD) have been associated with susceptibility to generalized epilepsy with febrile seizures [48]. Cholinergic receptor nicotinic alpha 3 subunit (CHRNA3) encodes a member of the nicotinic acetylcholine receptor family of proteins that likely plays a role in neurotransmission. Polymorphisms in this gene have been associated with an increased risk of smoking initiation and an increased susceptibility to lung cancer [49]. Glutamate ionotropic receptor AMPA type subunit 3 (GRIA3) which plays an important role in excitatory synaptic transmission is a candidate for bipolar disorder and nonspecific X-linked mental retardation [50]. GRM1 gene encodes a metabotropic glutamate receptor. It may be associated with many disease states, including schizophrenia, bipolar disorder, depression, and breast cancer [51]. Glutathione S-transferase pi 1 belongs to the group of enzymes that play important role in detoxification. GSTP1 variant proteins are thought to function in xenobiotic metabolism and play a role in susceptibility to cancer, and other diseases [52]. Endothelial PAS domain-containing protein 1 (EPAS1) also known as Hypoxia-inducible factor 2-alpha is a transcription factor which is induced as oxygen levels fall [53]. ATP2A2 catalyses the hydrolysis of ATP coupled with the translocation of calcium from the cytosol into the sarcoplasmic reticulum lumen and is involved in regulation of the contraction/relaxation cycle. Mutations in this gene cause Darier-White disease, also known as keratosis follicularis, an autosomal dominant skin disorder characterized by loss of adhesion between epidermal cells and abnormal keratinization. Other types of mutations in this gene have been associated with various forms of muscular dystrophies [54]. 5-Hydroxytryptamine Receptor 2B (HTR2B) is a serotonin receptor. It plays a role in the regulation of behaviour, including impulsive behaviour. It is required for normal proliferation of embryonic cardiac myocytes and normal heart development; protects cardiomyocytes against apoptosis and plays a role in the adaptation of pulmonary arteries to chronic hypoxia [55]. The Sox9 protein has been implicated in both initiation and progression of multiple solid tumors. Its role as a master regulator of morphogenesis during human development makes it an ideal candidate for perturbation in malignant tissues [56]. GRIN2C and GRIN2D are components of the NMDA receptors found in the central nervous system, are permeable to cations and have an important role in physiological processes such as learning, memory, and synaptic development. Alterations in the subunit composition of the receptor are associated with pathophysiological conditions such as Parkinson’s disease, Alzheimer’s disease, depression, and schizophrenia [57].

### 3.4 Interactions between genes downregulated in both neurotoxicity and cardiotoxicity

#### Gene regulatory interactions

The list of common downregulated genes between neurotoxicity and cardiotoxicity consisted only 2 genes-FLT4 and KDR, above the fold change cut off 3. When the list was subjected to gene regulatory interactions, seed node KDR along with intermediate nodes HHEX, HEY1, HIF2A/ARNT and ETS1 were included in the cluster (Fig. 10). KDR (VEGFR3) is a tyrosine-protein kinase that acts as a cell-surface receptor for VEGFA, VEGFC and VEGFD. Plays an essential role in the regulation of angiogenesis, vascular development, vascular permeability, and embryonic haematopoiesis[58].

**Figure 10.**
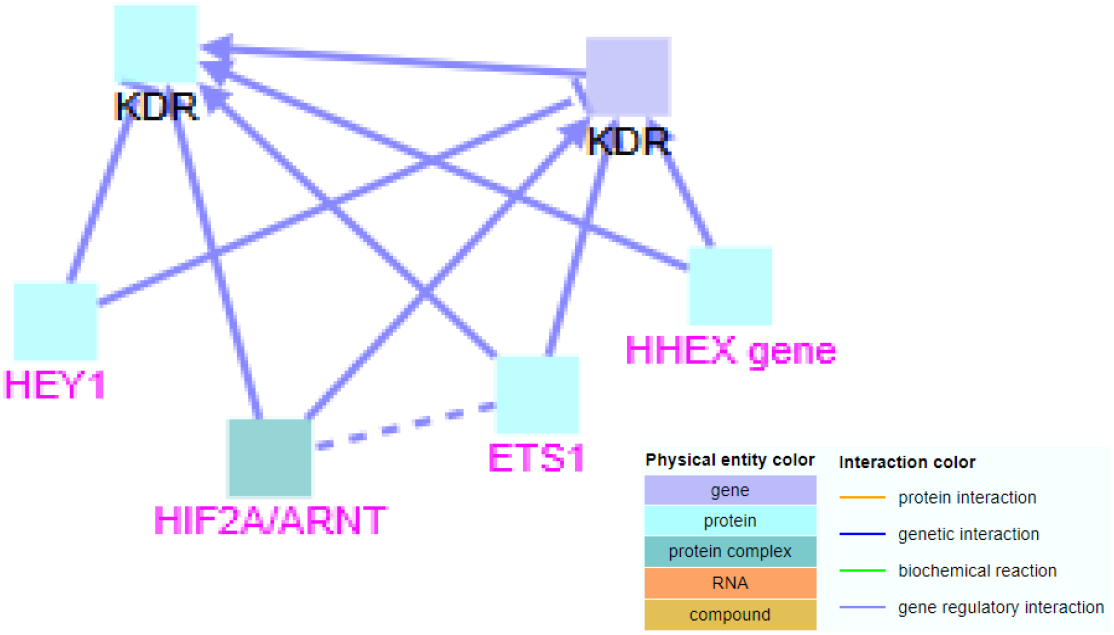
Gene regulatory interactions between genes downregulated both in neurotoxicity and cardiotoxicity. Black labels denote seed nodes and magenta labels denote intermediate nodes. Each edge represents an interaction.

#### Biochemical reactions

When biochemical reactions were included, the seed node FLT4 was included in the cluster (Fig. 11). FLT4 (VEGFR3) encodes a tyrosine kinase receptor for vascular endothelial growth factors C and D and plays an essential role in adult lymphangiogenesis and in the development of the vascular network and the cardiovascular system during embryonic development[59]. No genetic interactions were found.

**Figure 11.**
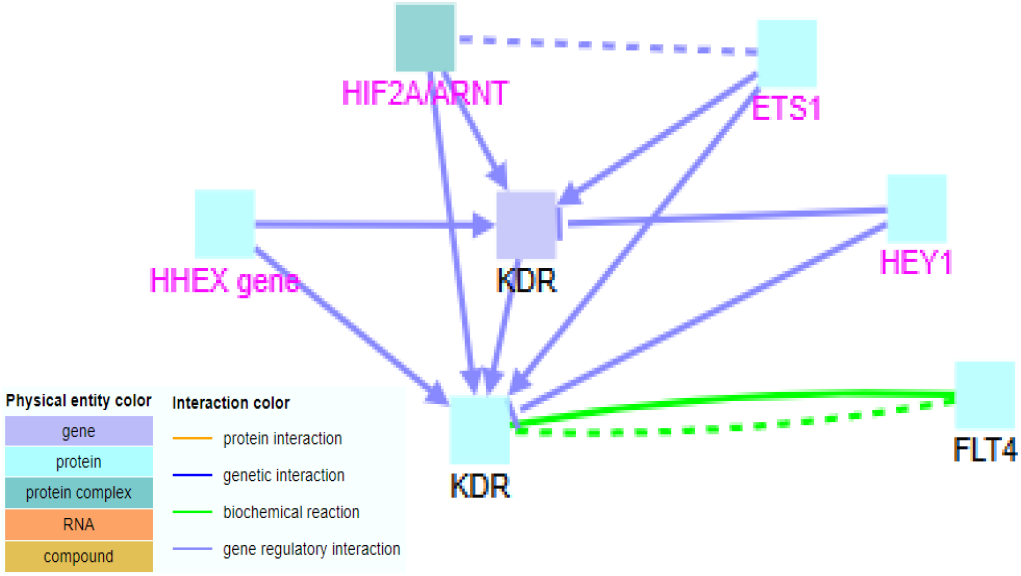
Gene regulatory interactions and biochemical reactions between genes downregulated both in neurotoxicity and cardiotoxicity. Black labels denote seed nodes and magenta labels denote intermediate nodes. Each edge represents an interaction.

#### Protein interactions

A protein interaction was found between the two seed nodes FLT4 and KDR was found (Fig. 12).

**Figure 12.**
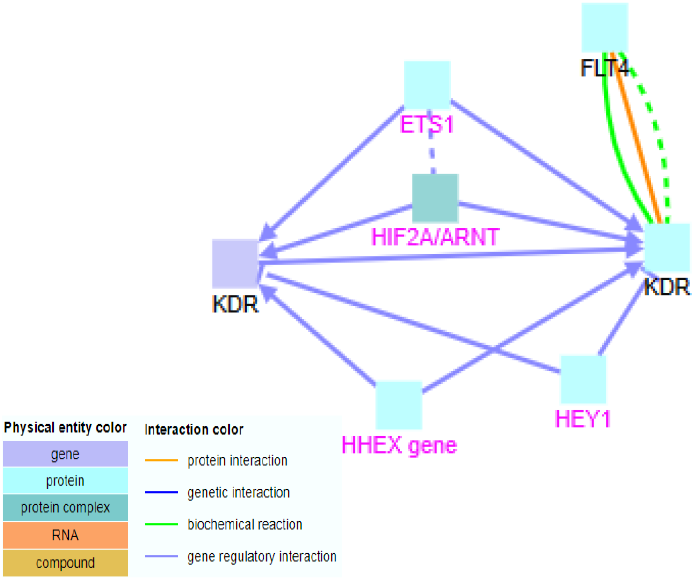
Gene regulatory, biochemical reactions and protein interactions between genes downregulated both in neurotoxicity and cardiotoxicity. Black labels denote seed nodes and magenta labels denote intermediate nodes. Each edge represents an interaction.

### 3.5 Interactions between genes upregulated exclusively in cardiotoxicity

#### Gene regulatory interactions

When the list containing upregulated genes in cardiotoxicity excluding the common upregulated genes in neurotoxicity and cardiotoxicity, were subjected to gene regulatory interactions, three clusters were formed (Fig. 13). Seed nodes like ESR1 in the first cluster, NR4A1 in the second cluster and IL1B in the third cluster, were included. The protein encoded by NR4A1 acts as a nuclear transcription factor. It is an orphan nuclear receptor. Translocation of the protein from the nucleus to mitochondria induces apoptosis [60]. The protein encoded by the IL1B gene is a member of the interleukin 1 cytokine family. This cytokine is an important mediator of the inflammatory response, and is involved in a variety of cellular activities, including cell proliferation, differentiation, and apoptosis [61]. Intermediate nodes like MTA1, MPG, MECP2, DDX17, JUN/FOS/ER alpha, ATF2 (dimer)/ER alpha and ATF2/JUN/ER alpha were included in the first cluster. Intermediate nodes like NFATc, MEF2D and MEF2D/NFAT1/Cbp/p300 were included in the second cluster. Intermediate nodes like STAT1-3 (dimer) and p-T180, Y182-MAPK11 were included in the third cluster.

**Figure 13.**
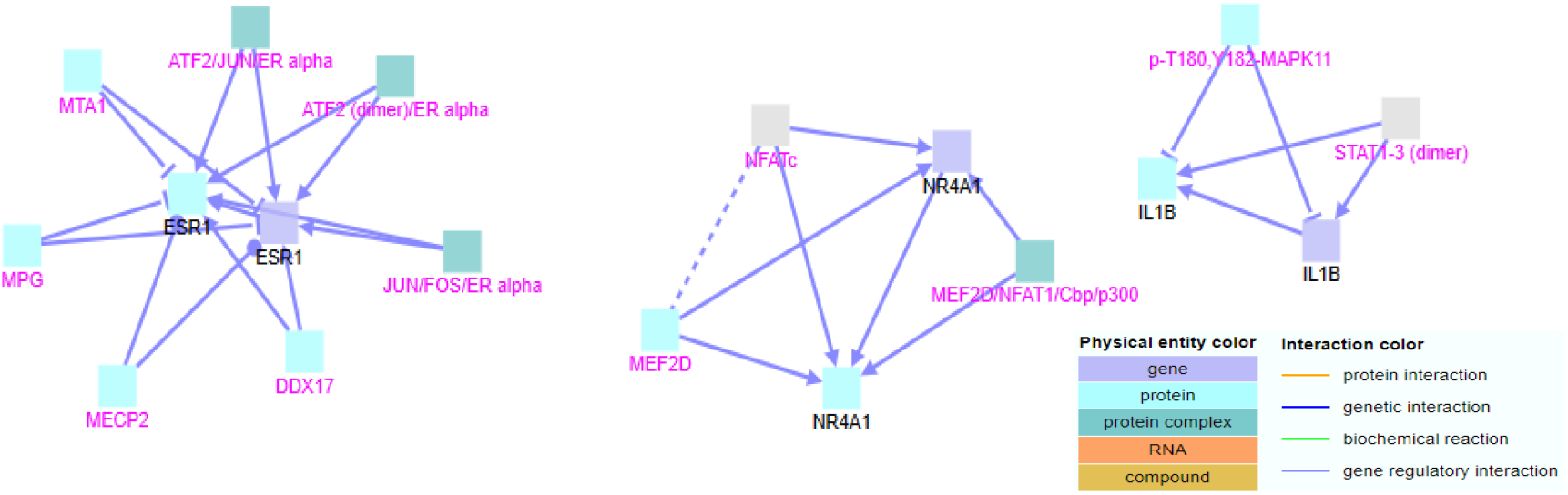
Gene regulatory interactions between genes upregulated exclusively in cardiotoxicity. Black labels denote seed nodes and magenta labels denote intermediate nodes. Each edge represents an interaction.

#### Biochemical reactions

When biochemical reactions were included in the analysis, the first and third cluster interacted and additional intermediate nodes like AP-1 complex, p300/CBP/RELA/p50, NFKB1, E2F5 and DNMT1 were found (Fig. 14). The second cluster previously formed remained the same here. A fourth cluster consisting of seed node CYP19A1 and additional intermediate nodes like JUND and FOXL2 was formed. also called estrogen synthetase, is a member of the cytochrome P450 superfamily The enzyme aromatase (CYP19A1) catalyses the conversion of androgen to estrogen, a rate-limiting step in estrogen biosynthesis [62].

**Figure 14.**
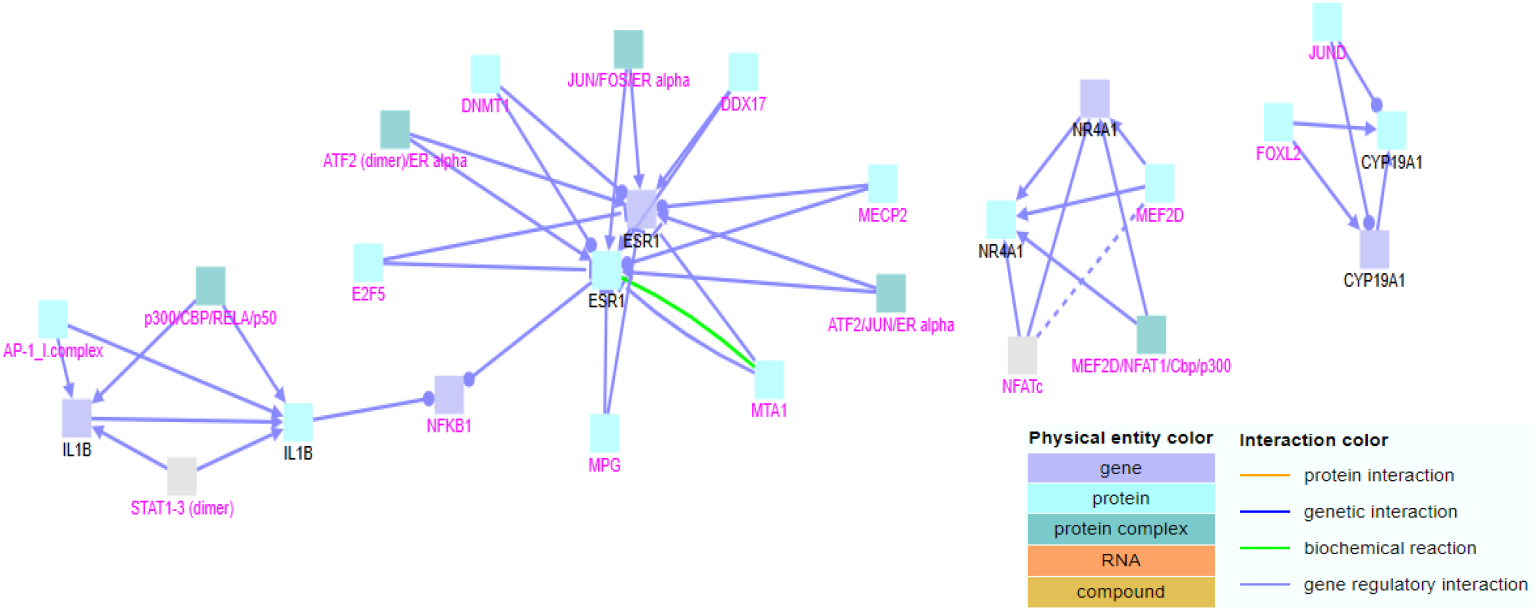
Gene regulatory interactions and biochemical reactions between genes upregulated exclusively in cardiotoxicity. Black labels denote seed nodes and magenta labels denote intermediate nodes. Each edge represents an interaction.

#### Genetic interactions

A genetic interaction between ESR1 and TFF1 gene was found (Fig. 15). Trefoil Factor 1 (TFF1) is a stable secretory protein expressed in gastrointestinal mucosa [63]. TFF1 expression is frequently lost in gastric carcinoma, probably through mechanism of DNA methylation, and it is therefore considered as a tumor suppressor gene [64]. Also strongly expressed in breast cancer [63].

**Figure 15.**
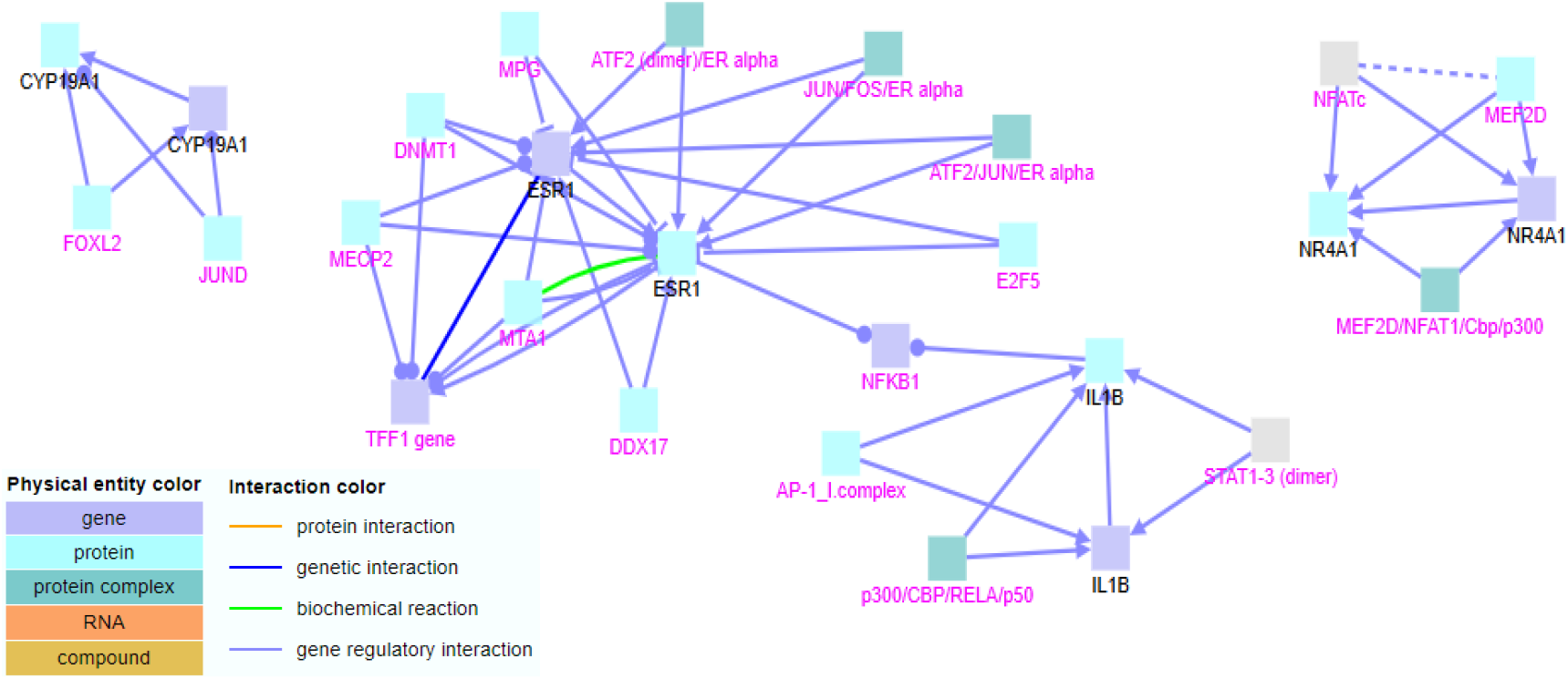
Gene regulatory, biochemical reactions and genetic interaction between genes upregulated exclusively in cardiotoxicity. Black labels denote seed nodes and magenta labels denote intermediate nodes. Each edge represents an interaction.

#### Protein interactions

When protein interactions were included in the analysis, all the clusters interacted (Fig. 16). Additional seed nodes like KAT6A and ARG2 and additional intermediate nodes like RLIM, CEBPD, TMOD3, VPRBP, AXIN2, E2, PELP1, ARG1, FOXO1, NCOA2 and NCOA3 were included. Chromosomal aberrations involving Histone acetyltransferase KAT6A may be a cause of acute myeloid leukaemia [65]. Arginase (ARG2) may play a role in the regulation of extra-urea cycle arginine metabolism and in down-regulation of nitric oxide synthesis. Arginine metabolism is a critical regulator of innate and adaptive immune responses. It is also involved in negative regulation of the survival capacity of activated CD4(+) and CD8(+) T cells [66].

**Figure 16.**
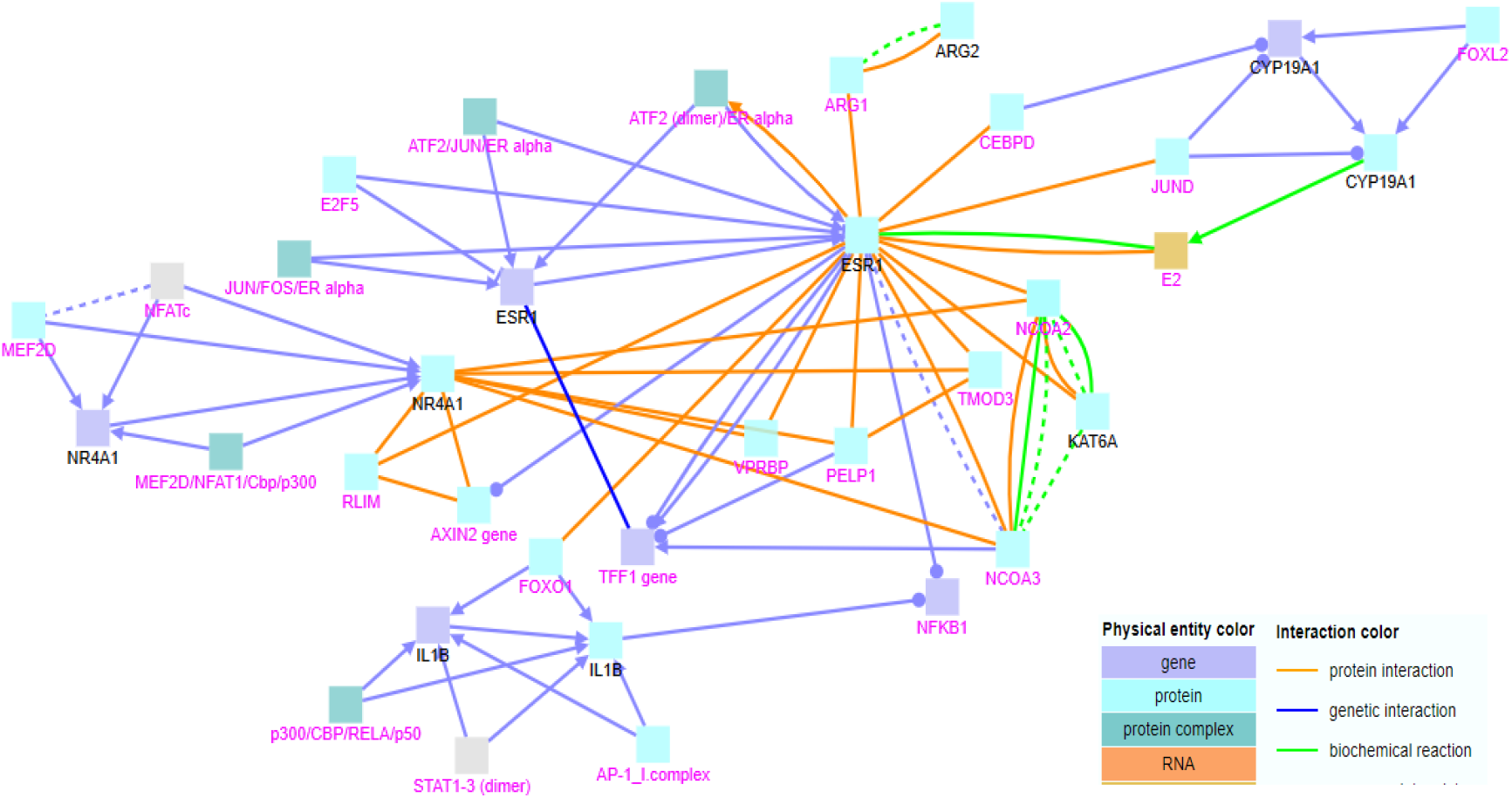
Gene regulatory, biochemical reactions, genetic and protein interactions between genes upregulated exclusively in cardiotoxicity._Black labels denote seed nodes and magenta labels denote intermediate nodes. Each edge represents an interaction.

### 3.6 Interactions between genes downregulated exclusively in cardiotoxicity

#### Gene regulatory interactions

When the list containing the downregulated genes in cardiotoxicity excluding the common downregulated genes in neurotoxicity and cardiotoxicity, was subjected to gene regulatory interactions, three clusters were formed (Fig. 17). Seed nodes like NOTCH1, VEGFA, TBX2 and intermediate nodes like PAS, VHL, JAG1 gene, TWIST1, HIF1A/ARNT/Cbp/p300/HDAC7 and SEMA4A:PLXND1 were included in the first cluster. Mutations in the NOTCH1 gene are associated with aortic valve disease, Adams-Oliver syndrome, T-cell acute lymphoblastic leukemia, chronic lymphocytic leukemia, and head and neck squamous cell carcinoma [67]. Vascular Endothelial Growth Factor A (VEGFA) is a growth factor that induces proliferation and migration of vascular endothelial cells and is essential for both physiological and pathological angiogenesis. This gene is upregulated in many known tumors and its expression is correlated with tumor stage and progression [68]. Studies suggest that the gene TBX2 may have a potential role in tumorigenesis as an immortalizing agent[69]. Seed node ITGB1 and additional intermediate node SRF (dimer)/MAL (dimer)/SCAI was included in the second cluster. Integrin Subunit Beta 1 (ITGB1) are membrane receptors involved in cell adhesion and recognition in a variety of processes including embryogenesis, homeostasis, tissue repair, immune response, and metastatic diffusion of tumor cells[70]. Seed node MEF2C and intermediate nodes SMAD2/SMAD2/SMAD4/FOXH1/NKX2-5 and CSX/GATA4 was included in the third cluster. This protein encoded by the Myocyte Enhancer Factor 2C (MEF2C) may play a role in maintaining the differentiated state of muscle cells. Mutations and deletions at this locus have been associated with severe cognitive disability, stereotypic movements, epilepsy, and cerebral malformation [71].

**Figure 17.**
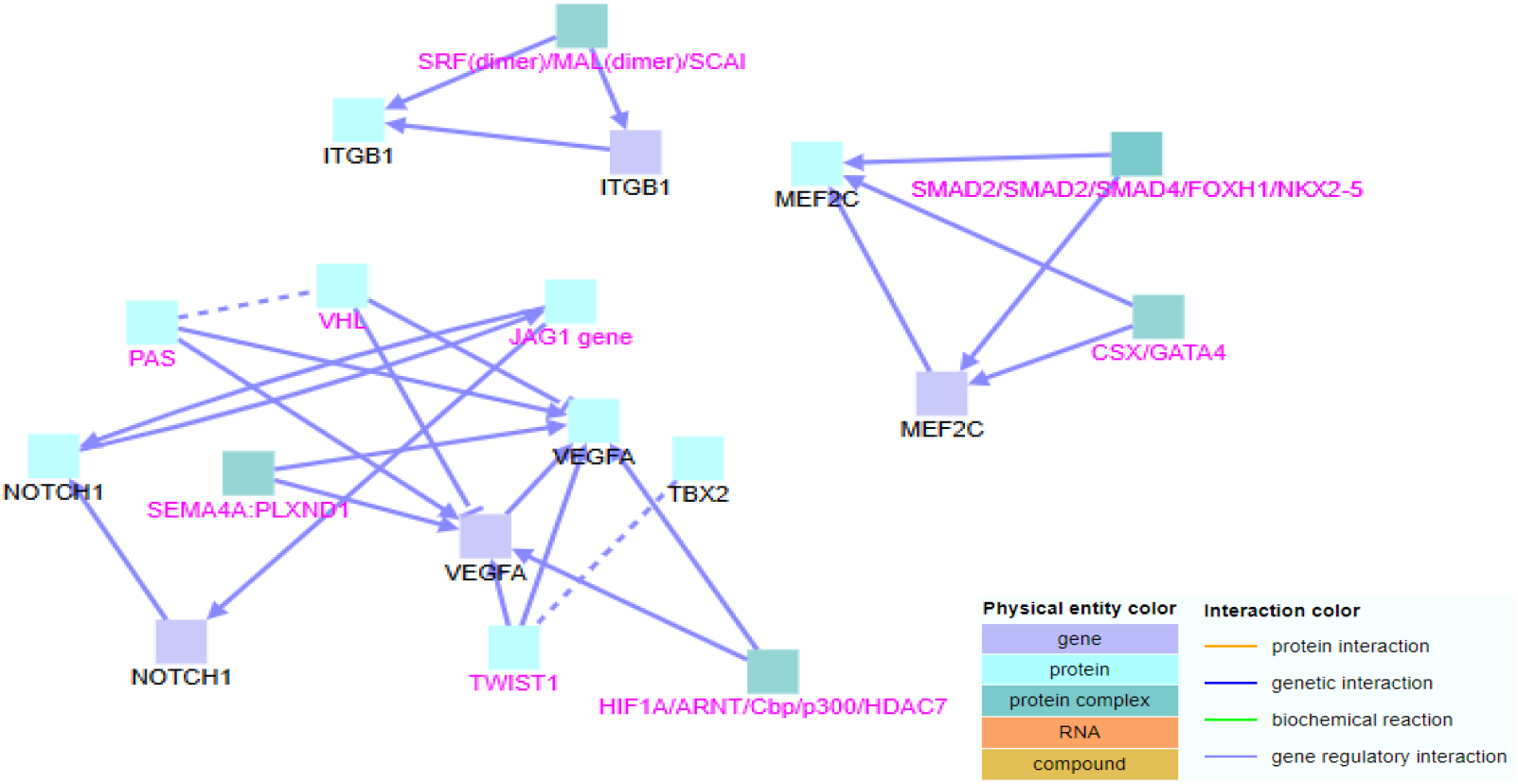
Gene regulatory interactions between genes downregulated exclusively in cardiotoxicity. Black labels denote seed nodes and magenta labels denote intermediate nodes. Each edge represents an interaction.

#### Biochemical interactions

When biochemical reactions were included, five clusters were formed (Fig. 18). Seed node VEGFA from the previously formed first cluster interacted with seed nodes ITGB1 and TBX2 here and additional intermediate nodes like HHEX gene, HIF-1 complex, HIF-1-alpha/ARNT/CREB/p300/JAB1/c-JUN, p-T611-FOXM1 and MYC/Max/MIZ-1 were included. The second cluster consisted of seed nodes NOTCH4 and NOTCH1 and intermediate nodes Gamma Secretase, gamma-secretase complex and JAG1. Mutations in the NOTCH4 gene may be associated with schizophrenia[72]. The third cluster previously formed remained the same. The fourth cluster consisted of seed nodes RHO, OPN1MW, OPN1SW and intermediate nodes like NFIC and 11cRAL. The protein encoded by the Rhodopsin (RHO) gene is found in rod cells in the back of the eye and is essential for vision in low-light conditions. Defects in this gene are a cause of congenital stationary night blindness [73]. The protein encoded by Opsin 1, Medium Wave Sensitive (OPN1MW) is called green cone photopigment or medium-wavelength sensitive opsin. Defects in this gene are the cause of deutanopic colour-blindness [74]. Opsin 1, Short Wave Sensitive (OPN1SW) encodes the blue cone pigment gene which is one of three types of cone photoreceptors responsible for normal color vision. Defects in this gene are the cause of tritan color blindness (tritanopia) [75]. The fifth cluster consisted of seed nodes PDE6C and PDE6H. The gene Phosphodiesterase 6C (PDE6C) encodes the alpha-prime subunit of cone phosphodiesterase. Mutations in this gene are associated with cone dystrophy type 4 (COD4) [76]. The gene Phosphodiesterase 6H (PDE6H) encodes the inhibitory (or gamma) subunit of the cone-specific cGMP phosphodiesterase. Mutations in this gene are associated with retinal cone dystrophy type 3A (RCD3A) [77].

**Figure 18.**
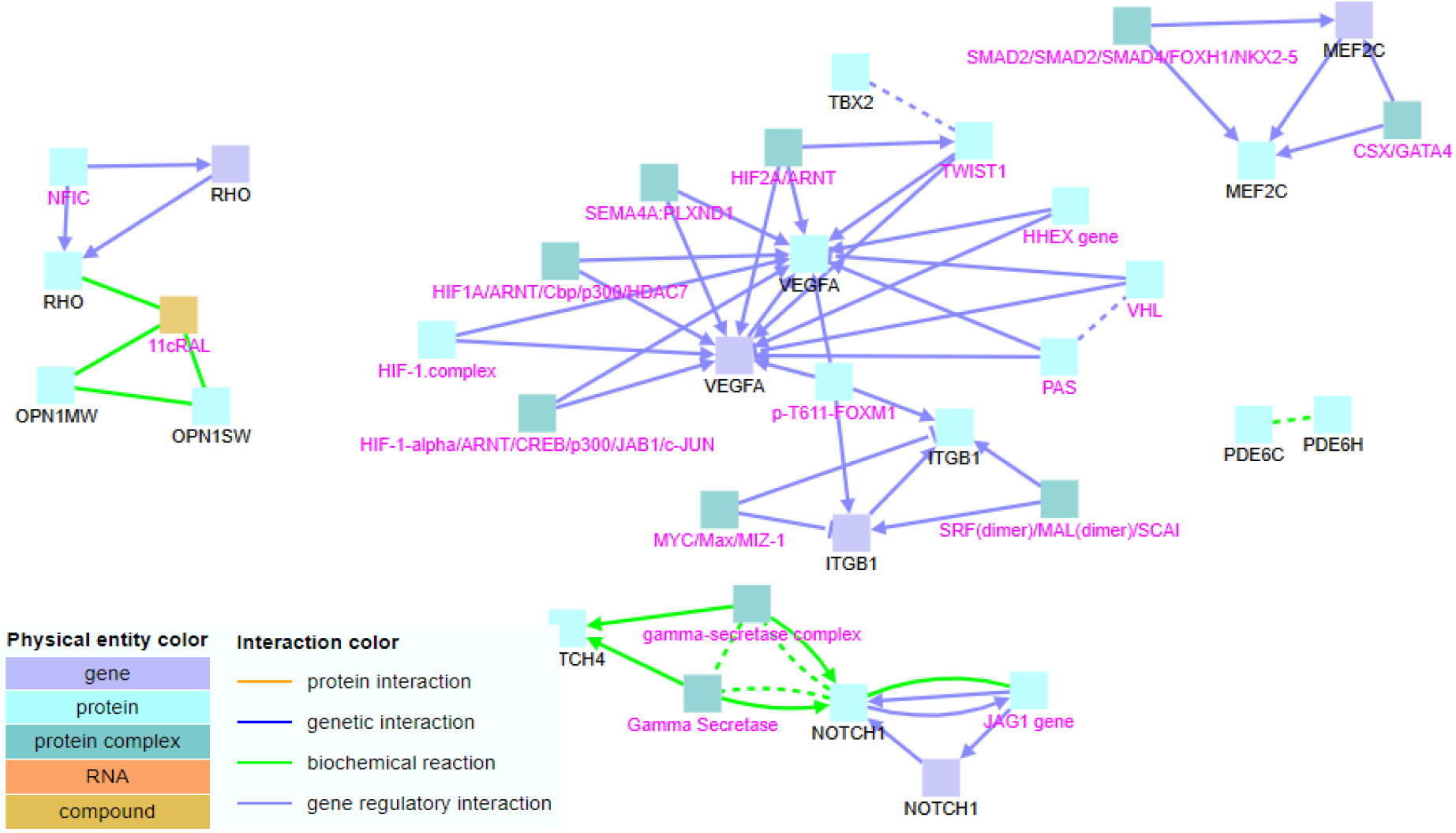
Gene regulatory interactions and biochemical reactions between genes downregulated exclusively in cardiotoxicity. Black labels denote seed nodes and magenta labels denote intermediate nodes. Each edge represents an interaction. No genetic interactions were found.

#### Protein interactions

When protein interactions were included in the analysis, the first three clusters interacted (Fig. 19). Additional seed nodes like SCN5A, NOTO, TBX6, CRX, TBXT and BMP2 were included. The protein encoded by the Sodium Voltage-Gated Channel Alpha Subunit 5 (SCN5A) gene is an integral membrane protein and tetrodotoxin-resistant voltage-gated sodium channel subunit. This protein is found primarily in cardiac muscle and is responsible for the initial upstroke of the action potential in an electrocardiogram. Defects in this gene are a cause of long QT syndrome type 3 (LQT3), an autosomal dominant cardiac disease [78]. The protein encoded by the gene Cone-Rod Homeobox (CRX) is a photoreceptor-specific transcription factor which plays a role in the differentiation of photoreceptor cells. This homeodomain protein is necessary for the maintenance of normal cone and rod function. Mutations in this gene are associated with photoreceptor degeneration [79]. BMP2 (Bone Morphogenetic Protein 2) is a Protein Coding gene. Diseases associated with Bone Morphogenetic Protein 2 (BMP2) include Short Stature, Facial Dysmorphism, And Skeletal Anomalies with or without Cardiac Anomalies 1 and Brachydactyly, Type A2 [80]. Previously formed fourth and fifth clusters remained the same.

**Figure 19.**
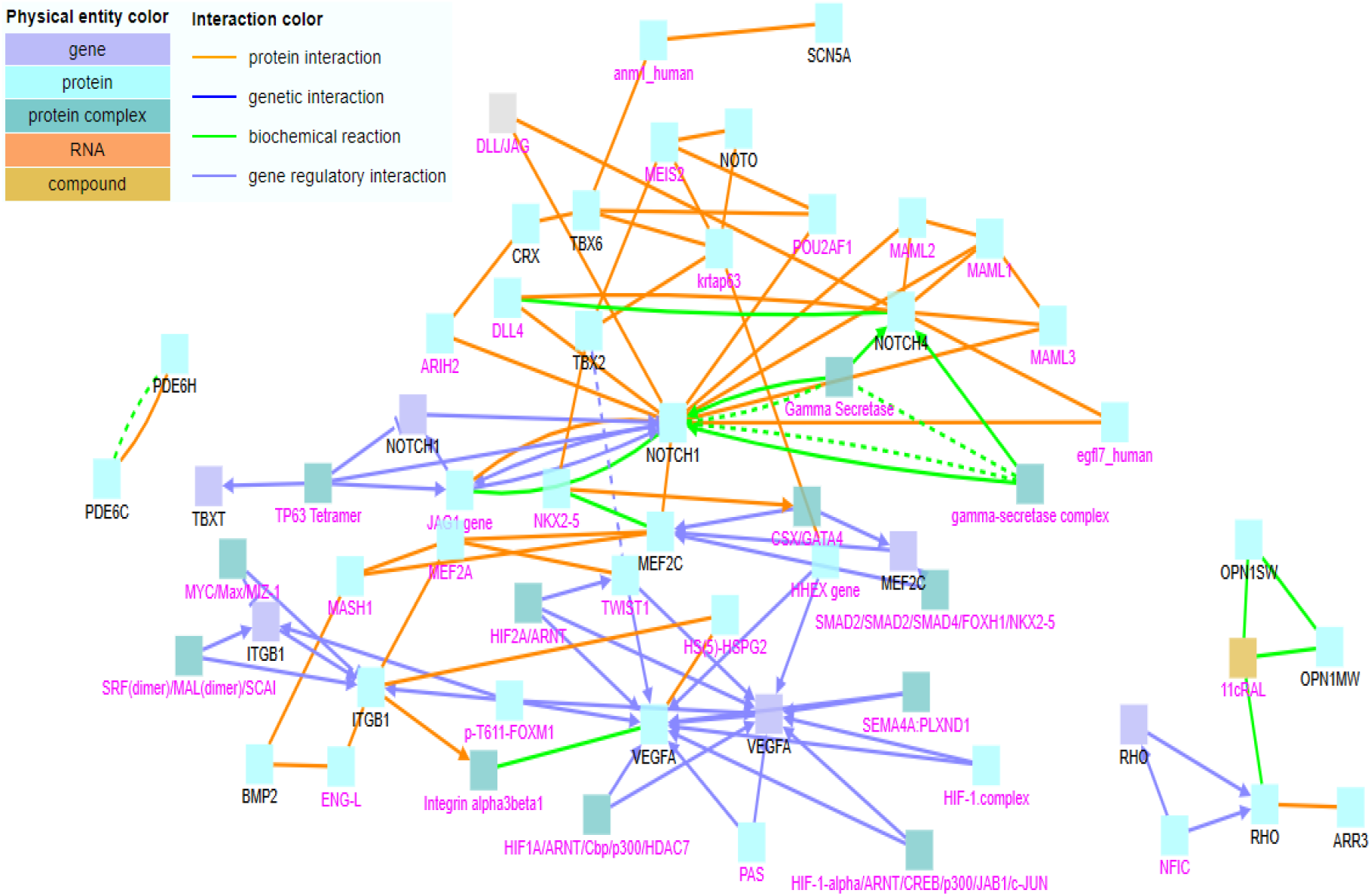
Gene regulatory, biochemical reactions and protein interactions between genes downregulated exclusively in cardiotoxicity. Black labels denote seed nodes and magenta labels denote intermediate nodes. Each edge represents an interaction.

## ACKNOWLEDGEMENTS

This work was supported by the Indian Institute of Hyderabad (IITH) and ECR-SERB-DST (ECR/2017/000242) grants to Dr Anamika Bhargava. Ministry of Education, India fellowship to Andrea Kagoo R and Department of Biotechnology, India fellowship to Ankush Sharma.

