## Supplementary information for "Interactions between genes altered during cardiotoxicity and neurotoxicity in zebrafish revealed using induced network modules analysis"

| TABLE S1: Articles that reported neurotoxicity | | |
|---|---|---|
| **S.no** | **Titles** | **DOI** |
| 1 | Naringenin alleviates 6-hydroxydopamine induced Parkinsonism in SHSY5Y cells and zebrafish model | 10.1016/j.cbpc.2020.108893 |
| 2 | Neurobehavioral effects of cyanobacterial biomass field extracts on zebrafish embryos and potential role of retinoids | 10.1016/j.aquatox.2020.105613 |
| 3 | An environmentally relevant mixture of polychlorinated biphenyls (PCBs) and polybrominated diphenylethers (PBDEs) disrupts mitochondrial function, lipid metabolism and neurotransmission in the brain of exposed zebrafish and their unexposed F2 offspring | 10.1016/j.scitotenv.2020.142097 |
| 4 | Developmental toxicity and neurotoxicity of penconazole enantiomers exposure on zebrafish (Danio rerio) | 10.1016/j.envpol.2020.115450 |
| 5 | Chronic exposure to bisphenol S induces oxidative stress, abnormal anxiety, and fear responses in adult zebrafish (Danio rerio) | 10.1016/j.scitotenv.2020.141633 |
| 6 | Parental exposure to environmental concentrations of tris(1,3-dichloro-2-propyl) phosphate induces abnormal DNA methylation and behavioral changes in F1 zebrafish larvae | 10.1016/j.envpol.2020.115305 |
| 7 | Integrated Hypoxia Signaling and Oxidative Stress in Developmental Neurotoxicity of Benzo[a]Pyrene in Zebrafish Embryos | 10.3390/antiox9080731 |
| 8 | Low-dose methylmercury exposure impairs the locomotor activity of zebrafish: Role of intestinal inositol metabolism | 10.1016/j.envres.2020.110020 |
| 9 | Embryonic atrazine exposure and later in life behavioral and brain transcriptomic, epigenetic, and pathological alterations in adult male zebrafish | 10.1007/s10565-020-09548-y |
| 10 | Sub-lethal toxicity assessment of the phenylurea herbicide linuron in developing zebrafish (Danio rerio) embryo/larvae | 10.1016/j.ntt.2020.106917 |

| | | |
|---|---|---|
| 11 | Toxicological effects induced on early life stages of zebrafish (Danio rerio) after an acute exposure to microplastics alone or co-exposed with copper | 10.1016/j.chemosphere.2020.127748 |
| 12 | Tributyltin enhanced anxiety of adult male zebrafish through elevating cortisol level and disruption in serotonin, dopamine and gamma-aminobutyric acid neurotransmitter pathways | 10.1016/j.ecoenv.2020.111014 |
| 13 | Protective Effects of Spermidine and Melatonin on Deltamethrin-Induced Cardiotoxicity and Neurotoxicity in Zebrafish | 10.1007/s12012-020-09591-5 |
| 14 | Effects of the chorion on the developmental toxicity of organophosphate esters in zebrafish embryos | 10.1016/j.jhazmat.2020.123389 |
| 15 | Diesel Exhaust Extract Exposure Induces Neuronal Toxicity by Disrupting Autophagy | 10.1093/toxsci/kfaa055 |
| 16 | Developmental and cardiac toxicities of propofol in zebrafish larvae | 10.1016/j.cbpc.2020.108838 |
| 17 | Low-Dose Exposure of Silica Nanoparticles Induces Neurotoxicity via Neuroactive Ligand-Receptor Interaction Signaling Pathway in Zebrafish Embryos | 10.2147/IJN.S254480 |
| 18 | Aluminum-Induced Cognitive Impairment and PI3K/Akt/mTOR Signaling Pathway Involvement in Occupational Aluminum Workers | 10.1007/s12640-020-00230-z |
| 19 | Effects of lincomycin hydrochloride on the neurotoxicity of zebrafish | 10.1016/j.ecoenv.2020.110725 |
| 20 | Effects of ecologically relevant concentrations of cadmium on locomotor activity and microbiota in zebrafish | 10.1016/j.chemosphere.2020.127220 |
| 21 | 8:8 Perfluoroalkyl phosphinic acid affects neurobehavioral development, thyroid disruption, and DNA methylation in developing zebrafish | 10.1016/j.scitotenv.2020.139600 |
| 22 | Calcium signaling as a possible mechanism behind increased locomotor response in zebrafish larvae exposed to a human relevant persistent organic pollutant mixture or PFOS | 10.1016/j.envres.2020.109702 |
| 23 | Dietary administration of probiotic Lactobacillus rhamnosus modulates the neurological toxicities of perfluorobutanesulfonate in zebrafish | 10.1016/j.envpol.2020.114832 |
| 24 | Nano-TiO(2) enhanced bioaccumulation and developmental neurotoxicity of bisphenol a in zebrafish larvae | 10.1016/j.envres.2020.109682 |
| 25 | Developmental exposure to mepanipyrim induces locomotor hyperactivity in zebrafish (Danio rerio) larvae | 10.1016/j.chemosphere.2020.127106 |
| 26 | Neuroprotective brain-derived neurotrophic factor signaling in the TAU-P301L tauopathy zebrafish model | 10.1016/j.phrs.2020.104865 |
| 27 | Differential responses of larval zebrafish to the fungicide propamocarb: Endpoints at development, locomotor behavior and oxidative stress | 10.1016/j.scitotenv.2020.139136 |

| | | |
|---|---|---|
| 28 | RNA-seq analysis and compound screening highlight multiple signalling pathways regulating secondary cell death after acute CNS injury in vivo | 10.1242/bio.050260 |
| 29 | Haloxyfop-P-methyl induces developmental defects in zebrafish embryos through oxidative stress and anti-vasculogenesis | 10.1016/j.cbpc.2020.108761 |
| 30 | Fenobucarb-induced developmental neurotoxicity and mechanisms in zebrafish | 10.1016/j.neuro.2020.03.013 |
| 31 | Titanium dioxide nanoparticles enhanced thyroid endocrine disruption of pentachlorophenol rather than neurobehavioral defects in zebrafish larvae | 10.1016/j.chemosphere.2020.126536 |
| 32 | Exposure to phthalates impaired neurodevelopment through estrogenic effects and induced DNA damage in neurons | 10.1016/j.aquatox.2020.105469 |
| 33 | Garcinol pacifies acrylamide induced cognitive impairments, neuroinflammation and neuronal apoptosis by modulating GSK signaling and activation of pCREB by regulating cathepsin B in the brain of zebrafish larvae | 10.1016/j.fct.2020.111246 |
| 34 | Environmental relevant concentrations of benzophenone-3 induced developmental neurotoxicity in zebrafish | 10.1016/j.scitotenv.2020.137686 |
| 35 | Corticotropin-releasing factor protects against ammonia neurotoxicity in isolated larval zebrafish brains | 10.1242/jeb.211540 |
| 36 | Triclosan induces zebrafish neurotoxicity by abnormal expression of miR-219 targeting oligodendrocyte differentiation of central nervous system | 10.1007/s00204-020-02661-1 |
| 37 | Tritiated Water Exposure in Zebrafish (Danio rerio): Effects on the Early-Life Stages | 10.1002/etc.4650 |
| 38 | Transcriptomic Responses of Bisphenol S Predict Involvement of Immune Function in the Cardiotoxicity of Early Life-Stage Zebrafish (Danio rerio) | 10.1021/acs.est.9b06213 |
| 39 | Impacts of Sex and Exposure Duration on Gene Expression in Zebrafish Following Perfluorooctane Sulfonate Exposure | 10.1002/etc.4628 |
| 40 | Inhibition of the electron transport chain in propofol induced neurotoxicity in zebrafish embryos | 10.1016/j.ntt.2020.106856 |
| 41 | Environmental co-exposure to TBT and Cd caused neurotoxicity and thyroid endocrine disruption in zebrafish, a three-generation study in a simulated environment | 10.1016/j.envpol.2019.113868 |
| 42 | Neuroprotective effects of mitoquinone and oleandrin on Parkinson's disease model in zebrafish | 10.1080/00207454.2019.1698567 |
| 43 | Bisphenol F-Induced Neurotoxicity toward Zebrafish Embryos | 10.1021/acs.est.9b04097 |
| 44 | The pyrethroid esfenvalerate induces hypoactivity and decreases dopamine transporter expression in embryonic/larval zebrafish (Danio rerio) | 10.1016/j.chemosphere.2019.125416 |
| 45 | Developmental exposure to lead at environmentally relevant concentrations impaired neurobehavior and NMDAR-dependent BDNF signaling in zebrafish larvae | 10.1016/j.envpol.2019.113627 |

| | | |
|---|---|---|
| 46 | Neurotoxic effects of aflatoxin B1 on human astrocytes in vitro and on glial cell development in zebrafish in vivo | 10.1016/j.jhazmat.2019.121639 |
| 47 | Environmentally relevant concentration of chromium induces nuclear deformities in erythrocytes and alters the expression of stress-responsive and apoptotic genes in brain of adult zebrafish | 10.1016/j.scitotenv.2019.135622 |
| 48 | Combined treatment of melatonin and sodium tanshinone IIA sulfonate reduced the neurological and cardiovascular toxicity induced by deltamethrin in zebrafish | 10.1016/j.chemosphere.2019.125373 |
| 49 | Bifenthrin induces developmental immunotoxicity and vascular malformation during zebrafish embryogenesis | 10.1016/j.cbpc.2019.108671 |
| 50 | Early-life exposure to the organophosphorus flame-retardant tris (1,3-dichloro-2-propyl) phosphate induces delayed neurotoxicity associated with DNA methylation in adult zebrafish | 10.1016/j.envint.2019.105293 |
| 51 | Fenvalerate triggers Parkinson-like symptom during zebrafish development through initiation of autophagy and p38 MAPK/mTOR signaling pathway | 10.1016/j.chemosphere.2019.125336 |
| 52 | Bisphenol F exposure impairs neurodevelopment in zebrafish larvae (Danio rerio) | 10.1016/j.ecoenv.2019.109870 |
| 53 | The food preservative ethoxyquin impairs zebrafish development, behavior and alters gene expression profile | 10.1016/j.fct.2019.110926 |
| 54 | The psychoactive cathinone derivative pyrovalerone alters locomotor activity and decreases dopamine receptor expression in zebrafish (Danio rerio) | 10.1002/brb3.1420 |
| 55 | Zebrafish behavioral phenomics employed for characterizing behavioral neurotoxicity caused by silica nanoparticles | 10.1016/j.chemosphere.2019.124937 |
| 56 | Clethodim exposure induced development toxicity and behaviour alteration in early stages of zebrafish life | 10.1016/j.envpol.2019.113218 |
| 57 | Ontogenetic expression of thyroid hormone signaling genes: An in vitro and in vivo species comparison | 10.1371/journal.pone.0221230 |
| 58 | Characterization of boscalid-induced oxidative stress and neurodevelopmental toxicity in zebrafish embryos | 10.1016/j.chemosphere.2019.124753 |
| 59 | Butylated Hydroxyanisole Exerts Neurotoxic Effects by Promoting Cytosolic Calcium Accumulation and Endoplasmic Reticulum Stress in Astrocytes | 10.1021/acs.jafc.9b02899 |
| 60 | Responses of pro- and anti-inflammatory cytokines in zebrafish liver exposed to sublethal doses of Aphanizomenon flosaquae DC-1 aphantoxins | 10.1016/j.aquatox.2019.105269 |
| 61 | Acute toxic effects of polyethylene microplastic on adult zebrafish | 10.1016/j.ecoenv.2019.109442 |
| 62 | Bioconcentration, depuration and toxicity of Pb in the presence of titanium dioxide nanoparticles in zebrafish larvae | 10.1016/j.aquatox.2019.105257 |
| 63 | A protective role of autophagy in Pb-induced developmental neurotoxicity in zebrafish | 10.1016/j.chemosphere.2019.06.227 |

| | | |
|---|---|---|
| 64 | The mechanisms underlying the developmental effects of bisphenol F on zebrafish | 10.1016/j.scitotenv.2019.05.489 |
| 65 | Neurodevelopmental toxicity assessments of alkyl phenanthrene and Dechlorane Plus co-exposure in zebrafish | 10.1016/j.ecoenv.2019.05.066 |
| 66 | Developmental neurotoxicity of reserpine exposure in zebrafish larvae (Danio rerio) | 10.1016/j.cbpc.2019.05.008 |
| 67 | Mitochondrial dysfunction-based cardiotoxicity and neurotoxicity induced by pyraclostrobin in zebrafish larvae | 10.1016/j.envpol.2019.04.122 |
| 68 | Comparative analyses of the neurobehavioral, molecular, and enzymatic effects of organophosphates on embryo-larval zebrafish (Danio rerio) | 10.1016/j.ntt.2019.04.002 |
| 69 | Identification of Potential Long Noncoding RNA Biomarker of Mercury Compounds in Zebrafish Embryos | 10.1021/acs.chemrestox.9b00029 |
| 70 | Exposure of low-dose fipronil enantioselectively induced anxiety-like behavior associated with DNA methylation changes in embryonic and larval zebrafish | 10.1016/j.envpol.2019.03.038 |
| 71 | Enantioselectivity of toxicological responses induced by maternal exposure of cis-bifenthrin enantiomers in zebrafish (Danio rerio) larvae | 10.1016/j.jhazmat.2019.03.049 |
| 72 | A simple method to study motor and non-motor behaviors in adult zebrafish | 10.1016/j.jneumeth.2019.03.008 |
| 73 | Zebrafish behavioral phenomics applied for phenotyping aquatic neurotoxicity induced by lead contaminants of environmentally relevant level | 10.1016/j.chemosphere.2019.02.174 |
| 74 | Effects of norfloxacin exposure on neurodevelopment of zebrafish (Danio rerio) embryos | 10.1016/j.neuro.2019.02.007 |
| 75 | Synergistic effects of Pb and repeated heat pulse on developmental neurotoxicity in zebrafish | 10.1016/j.ecoenv.2019.01.104 |
| 76 | Bioenergetic dysfunction in a zebrafish model of acute hyperammonemic decompensation | 10.1016/j.expneurol.2019.01.008 |

### TABLE S2: Articles that reported cardiotoxicity

| S.no | Titles | DOI |
|---|---|---|
| 1 | Famoxadone-cymoxanil induced cardiotoxicity in zebrafish embryos | 10.1016/j.ecoenv.2020.111339 |
| 2 | Development toxicity and cardiotoxicity in zebrafish from exposure to iprodione | 10.1016/j.chemosphere.2020.127860 |
| 3 | Acute fluorene-9-bisphenol exposure damages early development and induces cardiotoxicity in zebrafish (Danio rerio) | 10.1016/j.ecoenv.2020.110922 |
| 4 | Protective Effects of Spermidine and Melatonin on Deltamethrin-Induced Cardiotoxicity and Neurotoxicity in Zebrafish | 10.1007/s12012-020-09591-5 |

| | | |
|---|---|---|
| 5 | Potential Molecular Mechanisms and Drugs for Aconitine-Induced Cardiotoxicity in Zebrafish through RNA Sequencing and Bioinformatics Analysis | 10.12659/MSM.924092 |
| 6 | Risk assessment of cardiotoxicity to zebrafish (Danio rerio) by environmental exposure to triclosan and its derivatives | 10.1016/j.envpol.2020.114995 |
| 7 | Exposure to Oxadiazon-Butachlor causes cardiac toxicity in zebrafish embryos | 10.1016/j.envpol.2020.114775 |
| 8 | Resveratrol protects against PM2.5-induced heart defects in zebrafish embryos as an antioxidant rather than as an AHR antagonist | 10.1016/j.taap.2020.115029 |
| 9 | Oxidative stress in bisphenol AF-induced cardiotoxicity in zebrafish and the protective role of N-acetyl N-cysteine | 10.1016/j.scitotenv.2020.139190 |
| 10 | A Phenotypic and Genotypic Evaluation of Developmental Toxicity of Polyhexamethylene Guanidine Phosphate Using Zebrafish Embryo/Larvae | 10.3390/toxics8020033 |
| 11 | Exposure to pyrimethanil induces developmental toxicity and cardiotoxicity in zebrafish | 10.1016/j.chemosphere.2020.126889 |
| 12 | Downregulation of miR-133a contributes to the cardiac developmental toxicity of trichloroethylene in zebrafish | 10.1016/j.chemosphere.2020.126610 |
| 13 | Isoniazid causes heart looping disorder in zebrafish embryos by the induction of oxidative stress | 10.1186/s40360-020-0399-2 |
| 14 | Exposure to Crude Oil Induces Retinal Apoptosis and Impairs Visual Function in Fish | 10.1021/acs.est.9b07658 |
| 15 | Chiral toxicity of muscone to embryonic zebrafish heart | 10.1016/j.aquatox.2020.105451 |
| 16 | α-asarone induces cardiac defects and QT prolongation through mitochondrial apoptosis pathway in zebrafish | 10.1016/j.toxlet.2020.02.003 |
| 17 | Paeonol Reverses Adriamycin Induced Cardiac Pathological Remodeling through Notch1 Signaling Reactivation in H9c2 Cells and Adult Zebrafish Heart | 10.1021/acs.chemrestox.9b00093 |
| 18 | Retinoid X receptor alpha is a spatiotemporally predominant therapeutic target for anthracycline-induced cardiotoxicity | 10.1126/sciadv.aay2939 |

| | | |
|---|---|---|
| 19 | Aconitine induces cardiotoxicity through regulation of calcium signaling pathway in zebrafish embryos and in H9c2 cells | 10.1002/jat.3943 |
| 20 | Cardiotoxicity and Cardioprotection by Artesunate in Larval Zebrafish | 10.1177/1559325819897180 |
| 21 | Fucoidan Derived from Fucus vesiculosus Inhibits the Development of Human Ovarian Cancer via the Disturbance of Calcium Homeostasis, Endoplasmic Reticulum Stress, and Angiogenesis | 10.3390/md18010045 |
| 22 | Transcriptomic Responses of Bisphenol S Predict Involvement of Immune Function in the Cardiotoxicity of Early Life-Stage Zebrafish (Danio rerio) | 10.1021/acs.est.9b06213 |
| 23 | Exposure to diclofop-methyl induces cardiac developmental toxicity in zebrafish embryos | 10.1016/j.envpol.2020.113926 |
| 24 | AHR-mediated ROS production contributes to the cardiac developmental toxicity of PM2.5 in zebrafish embryos | 10.1016/j.scitotenv.2019.135097 |
| 25 | Combined treatment of melatonin and sodium tanshinone IIA sulfonate reduced the neurological and cardiovascular toxicity induced by deltamethrin in zebrafish | 10.1016/j.chemosphere.2019.125373 |
| 26 | Induction of developmental toxicity and cardiotoxicity in zebrafish embryos/larvae by acetyl-11-keto-β-boswellic acid (AKBA) through oxidative stress | 10.1080/01480545.2019.1663865 |
| 27 | Exposure to water-accommodated fractions of two different crude oils alters morphology, cardiac function and swim bladder development in early-life stages of zebrafish | 10.1016/j.chemosphere.2019.06.199 |
| 28 | Cardiotoxicity of forchlorfenuron (CPPU) in zebrafish (Danio rerio) and H9c2 cardiomyocytes | 10.1016/j.chemosphere.2019.06.027 |
| 29 | Mitochondrial dysfunction-based cardiotoxicity and neurotoxicity induced by pyraclostrobin in zebrafish larvae | 10.1016/j.envpol.2019.04.122 |
| 30 | Glyphosate induces toxicity and modulates calcium and NO signaling in zebrafish embryos | 10.1016/j.bbrc.2019.04.074 |

| | | |
|---|---|---|
| 31 | Cardiovascular Effects of PCB 126 (3,3',4,4',5-Pentachlorobiphenyl) in Zebrafish Embryos and Impact of Co-Exposure to Redox Modulating Chemicals | 10.3390/ijms20051065 |
| 32 | Developmental toxicity of triclocarban in zebrafish (Danio rerio) embryos | 10.1002/jbt.22289 |
| 33 | Aconitum alkaloids induce cardiotoxicity and apoptosis in embryonic zebrafish by influencing the expression of cardiovascular relative genes | 10.1016/j.toxlet.2019.01.002 |
| 34 | Cardiogenesis impairment promoted by bisphenol A exposure is successfully counteracted by epigallocatechin gallate | 10.1016/j.envpol.2019.01.004 |

| **TABLE S3: Exclusively upregulated genes in neurotoxicity** | |
|---|---|
| **Gene name** | **Fold Change values relative to control** |
| *LTA* | 16 |
| *malat1* | 10.79 |
| *SLC7A5* | 10 |
| *klf9* | 10 |
| *CXCL8* | 9.6 |
| *IL10* | 7.85 |
| *tnfsf11* | 7.5 |
| *thra* | 7 |
| *guca1c* | 6.44 |
| *HMOX1* | 6 |
| *tuba1b* | 6 |
| *odc1* | 5 |
| *dio3* | 5 |
| *ambra1* | 4.5 |
| *PRKAB1* | 4 |
| *smox* | 4 |
| *NFE2L2* | 3.78 |
| *TGFB3* | 3.52 |
| *tp53* | 3.5 |
| Total | 19 |
| **GENES NOT INCLUDED** | |
| *Il6* | 3 |
| *ulk1* | 3 |
| *pmel* | 3 |
| *tshr* | 2.9 |
| *mt2a* | 2.72 |

| | |
|---|---|
| *bax* | 2.5 |
| *NKX2-1* | 2.5 |
| *crh* | 2.4 |
| *dio3* | 2.4 |
| *casp3* | 2.2 |
| *ulk1* | 2.1 |
| *atg7* | 2 |
| *POMC* | 2 |
| *sox2* | 2 |
| *PDGFRA* | 2 |
| *ephA4* | 2 |
| *thra* | 2 |
| *HSP90AA1* | 2 |
| *GPT2* | 2 |
| *dnmt3a* | 1.98 |
| *oca2* | 1.8 |
| *casp8* | 1.75 |
| *BICDL1* | 1.75 |
| *Hoxb1* | 1.75 |
| *rxrg* | 1.6 |
| *keap1* | 1.5 |
| *hmox1* | 1.5 |
| *ahr* | 1.5 |
| *dnmt3b* | 1.46 |
| *lyz* | 1.45 |
| *casp3* | 1.4 |
| *MAP2* | 1.375 |
| *CRHR2* | 1.25 |
| *crh* | 1.2 |
| *PCCB* | 1.2 |
| *slc27a1* | 1.16 |
| *SLC16A2* | 1 |
| *SLCO1C1* | 1 |
| *Eno2* | 1 |
| *PCCA* | 1 |
| Total | 40 |

**TABLE S4: Common upregulated genes in neurotoxicity and cardiotoxicity**

| Gene name | Fold Change values relative to control |
|---|---|
| *cyp1a1* | 42.74 |
| *ptgs2* | 12.27 |
| *hipk2* | 4.5 |
| *il21r* | 3.1 |

| | |
|---|---|
| *jam2* | 3.1 |
| *aqp1* | 3.1 |
| Total | 6 |
| **GENES NOT INCLUDED** | |
| *vipr2* | 3 |
| *hnf1a* | 3 |
| *il12b* | 2.8 |
| *ANGPT2* | 2.7 |
| *slc7a11* | 2.5 |
| *csf3* | 2.4 |
| *TLR4* | 2 |
| *nos2* | 2 |
| *wnt6* | 2 |
| *SULT1A1* | 1.74 |
| *agxt* | 1.67 |
| *vtg1* | 1.18 |
| Total | 12 |

| **TABLE S5: Exclusively downregulated genes in neurotoxicity** | |
|---|---|
| **Gene name** | **Fold Change values relative to control (negative values)** |
| *gstp1* | 50 |
| *grm1* | 20 |
| *matn3* | 11.11111111 |
| *nlgn2* | 11.11111111 |
| *mmp9* | 10 |
| *grm6* | 10 |
| *chrna3* | 10 |
| *esr1* | 10 |
| *tfap2e* | 10 |
| *atp2a2* | 10 |
| *ryr1* | 6.666666667 |
| *cacna1f* | 6.493506494 |
| *Arrb2* | 6.211180124 |
| *cacna1a* | 6.211180124 |
| *lhcgr* | 5.917159763 |
| *cacna1i* | 5.917159763 |
| *egr2* | 5.882352941 |
| *gria3* | 5.555555556 |
| *EPAS1* | 5 |
| *npffr2* | 5 |
| *gabrg3* | 5 |

| | |
|---|---|
| *glra3* | 5 |
| *MTOR* | 5 |
| *cdk6* | 5 |
| *ryr1* | 5 |
| *socs3* | 4.761904762 |
| *cdc42* | 4.761904762 |
| *cacna1d* | 4.484304933 |
| *grin2d* | 4.484304933 |
| *grin2c* | 4.484304933 |
| *igf1* | 4 |
| *PIK3CA* | 4 |
| *SLC6A1* | 4 |
| *gria2* | 4 |
| *fos* | 4 |
| *gstp1* | 4 |
| *htr2b* | 3.90625 |
| *Htr5a* | 3.333333333 |
| *sox9* | 3.333333333 |
| *gabrd* | 3.333333333 |
| Total | 40 |
| **GENES NOT INCLUDED** | |
| *igfbp1* | 2.564102564 |
| *th* | 2.55 |
| *kcnn2* | 2.5 |
| *ric3* | 2.5 |
| *grin2a* | 2.5 |
| *AKT1* | 2.5 |
| *gls* | 2.5 |
| *maob* | 2.5 |
| *dio1* | 2.5 |
| *cdk2* | 2.5 |
| *adh5* | 2.325581395 |
| *cat* | 2.222222222 |
| *hif1a* | 2.222222222 |
| *Col8a1* | 2.222222222 |
| *tap1* | 2.222222222 |
| *zc4h2* | 2.16 |
| *pax6* | 2.083333333 |
| *gria4* | 2.083333333 |
| *HTR1A* | 2 |
| *Htr1b* | 2 |
| *dnmt3a* | 2 |
| *drd2* | 2 |
| *gabrg2* | 2 |

| | |
|---|---|
| *fev* | 2 |
| *pomc* | 2 |
| *chrnb3* | 2 |
| *npy4r* | 2 |
| *grin1* | 2 |
| *olig2* | 2 |
| *myod1* | 2 |
| *neurod1* | 2 |
| *Mog* | 2 |
| *ccnd1* | 2 |
| *sod2* | 2 |
| *gabra1* | 2 |
| *tfap2a* | 2 |
| *wnt3a* | 2 |
| *wnt5a* | 2 |
| *drd4* | 1.89 |
| *pi4k2a* | 1.818181818 |
| *cyc1* | 1.818181818 |
| *Nr4a2* | 1.666666667 |
| *shh* | 1.666666667 |
| *adra2b* | 1.666666667 |
| *Mag* | 1.666666667 |
| *ttr* | 1.666666667 |
| *grin2c* | 1.666666667 |
| *ccne1* | 1.666666667 |
| *gadd45a* | 1.666666667 |
| *hdc* | 1.5625 |
| *slc1a2* | 1.538461538 |
| *otp* | 1.538461538 |
| *drd2c* | 1.46 |
| *chrm5* | 1.428571429 |
| *slc6a4* | 1.428571429 |
| *ntrk2* | 1.428571429 |
| *Th* | 1.333333333 |
| *th* | 1.333333333 |
| *scn1b* | 1.333333333 |
| *il1rapl1* | 1.333333333 |
| *brinp3* | 1.333333333 |
| *drd2* | 1.333333333 |
| *gad2* | 1.333333333 |
| *crhbp* | 1.333333333 |
| *tuba1a* | 1.333333333 |
| *Plp1* | 1.333333333 |
| *DIABLO* | 1.333333333 |
| *rad51* | 1.333333333 |
| *otp* | 1.333333333 |

| | |
|---|---|
| *cep135* | 1.265822785 |
| *slc6a3* | 1.25 |
| *acth* | 1.25 |
| *acsl4* | 1.219512195 |
| *insig1* | 1.219512195 |
| *DNMT3L* | 1.162790698 |
| *HTR1A* | 1.111111111 |
| *Htr2a* | 1.111111111 |
| *Elavl3* | 1.111111111 |
| Total | 78 |

| **TABLE S6: Common downregulated genes in neurotoxicity and cardiotoxicity** | |
|---|---|
| **Gene name** | **Fold Change values relative to control (negative values)** |
| *flt4* | 5 |
| *kdr* | 3.333333333 |
| Total | 2 |
| **GENES NOT INCLUDED** | |
| *vegfc* | 2.857142857 |
| *klf2* | 2.702702703 |
| *sod1* | 2.5 |
| *wnt8a* | 1.886792453 |
| *cox8a* | 1.785714286 |
| *flt1* | 1.666666667 |
| *UQCRC2* | 1.612903226 |
| *ATP5MC3* | 1.587301587 |
| Total | 8 |

| **TABLE S7: Exclusively upregulated genes in cardiotoxicity** | |
|---|---|
| **Gene name** | **Fold Change values relative to control** |
| *IL1B* | 17.1 |
| *CYP19A1* | 12.9 |
| *Esr1* | 6 |
| *nr4a1* | 5 |
| *Arg2* | 3.61 |
| *kat6a* | 3.5 |
| Total | 6 |
| **GENES NOT INCLUDED** | |

| | |
|---|---|
| *serpine1* | 3 |
| *ugdh* | 2.6 |
| *pcna* | 2.6 |
| *hspb11* | 2.5 |
| *tbx5* | 2.1 |
| *COX4I1* | 1.88 |
| *hspA4* | 1.72 |
| *nqo1* | 1.5 |
| Total | 8 |

| **TABLE S8: Exclusively downregulated genes in cardiotoxicity** | |
|---|---|
| **Gene name** | **Fold Change values relative to control (negative values)** |
| *OPN1MW* | 50.00 |
| *Rgr* | 50.00 |
| *OPN1SW* | 50.00 |
| *Crx* | 40.00 |
| *Pde6c* | 40.00 |
| *Pde6h* | 33.33 |
| *Rho* | 33.33 |
| *scn5a* | 20.00 |
| *Arr3* | 20.00 |
| *Gnat2* | 20.00 |
| *Rpe65* | 20.00 |
| *notch1* | 8.33 |
| *tbxt* | 5.00 |
| *noto* | 5.00 |
| *tbx6* | 5.00 |
| *Trh* | 5.00 |
| *vegfa* | 5.00 |
| *Tbx2* | 4.00 |
| *Bmp2* | 4.00 |
| *notch4* | 4.00 |
| *mef2c* | 4.00 |
| *ITGB1* | 3.33 |
| Total | 22 |
| **GENES NOT INCLUDED** | |
| *Arg1* | 2.78 |
| *cacng6* | 2.78 |
| *TNC* | 2.50 |
| *Hey2* | 2.50 |

| | |
|---|---|
| *casp8* | 2.50 |
| *cacnb3* | 2.50 |
| *mga* | 2.27 |
| *tbx5* | 2.22 |
| *axin2* | 2.22 |
| *bmp4* | 2.00 |
| *ctnnb1* | 2.00 |
| *Lef1* | 2.00 |
| *TNNT2* | 2.00 |
| *Thrb* | 2.00 |
| *gata1* | 2.00 |
| *Tbx1* | 2 |
| *camk2a* | 1.87 |
| *cacna2d4* | 1.85 |
| *notch1* | 1.82 |
| *cacna1a* | 1.67 |
| *gpx1* | 1.54 |
| *myh7* | 1.45 |
| *ppp2r3a* | 1.42 |
| *atp2a1* | 1.33 |
| *tanc1* | 1.33 |
| *ryr2* | 0.5 |
| *cacna1c* | 0.1 |
| Total | 27 |
